# Nucleus accumbens D1-receptors regulate and focus transitions to reward-seeking action

**DOI:** 10.1101/2021.06.15.448563

**Authors:** LL Grima, MC Panayi, O Haermson, E Syed, SG Manohar, M Husain, ME Walton

**Author notes:** Correspondence: Laura Grima, Mark Walton.

## Abstract

While it is well established that dopamine transmission is integral in mediating the influence of reward expectations on reward-seeking actions, the precise causal role of dopamine transmission in moment-to-moment cue-driven behavioural control remains contentious. This is a particular issue in situations where it is necessary to refrain from responding to achieve a beneficial outcome. To examine this, we manipulated dopamine transmission pharmacologically as rats performed a Go/No-Go task that required them to either make or withhold action to gain either a small or large reward. Stimulation of D1Rs, both globally and locally in the nucleus accumbens core (NAcC) region consistently disrupted No-Go performance, potentiating inappropriate responses that clustered strongly just after cue presentation. D1R blockade did not, however, improve rats’ ability to withhold responses, but instead primarily disrupted performance on Go trials. While global D1R blockade caused a general reduction of invigoration of reward seeking actions, intra-NAcC administration of the D1R antagonist by contrast increased the likelihood that Go trial performance was in an “unfocused” state. Such a state was characterised, both on and off drug, by a reduction in the precision and speed of responding even though the appropriate action sequence was often executed. These findings suggests that the balance of activity at NAcC D1Rs plays a key role in enabling the rapid activation of a focused, reward-seeking state to enable animals to efficiently and accurately achieve their goal.

## Introduction

The balance of dopamine transmission plays a key role in mediating the efficacy of reward-guided behaviour (Dalley & Roiser, 2012; Floresco, 2015; Nicola, 2010; Robbins & Everitt, 2007). Reduction of dopamine transmission in ventral striatal regions such as the nucleus accumbens core (NAcC) reduces the likelihood of responding to reward-associated cues and disrupts the willingness to persist with instrumental responses (du Hoffmann & Nicola, 2014; Salamone, Correa, Farrar, & Mingote, 2007; Yun, Nicola, & Fields, 2004). Conversely, hyperdopaminergic states can also result in dysfunctional reward pursuit (Murphy, Robinson, Theobald, Dalley, & Robbins, 2008; Pattij, Janssen, Vanderschuren, Schoffelmeer, & Van Gaalen, 2007; Pezze, Dalley, & Robbins, 2007; Van Gaalen, Brueggeman, Bronius, Schoffelmeer, & Vanderschuren, 2006).

However, the precise relationship between reward expectation, dopamine transmission and behavioural control remains unclear. It is well established that the presentation of reward-associated cues rapidly causes changes in dopamine activity, the magnitude of which reflects the subjective value of the expected future reward (Collins, Aitken, Greenfield, Ostlund, & Wassum, 2016; Gan, Walton, & Phillips, 2010; Lak, Stauffer, & Schultz, 2014; Papageorgiou, Baudonnat, Cucca, & Walton, 2016). This acts to influence the activity of striatal medium spiny neurons (MSNs), particularly those expressing D1-like receptors (D1Rs) (Dreher & Jackson, 1989; Lahiri & Bevan, 2020; Nicola, Taha, Kim, & Fields, 2005; Oldenburg & Sabatini, 2015; Richfield, Penney, & Young, 1989; Tritsch & Sabatini, 2012).

Different theories of dopamine posit that, on the one hand, it facilitates action *per se* through increasing vigour (Dayan, 2012; Niv, Daw, Joel, & Dayan, 2007) or on the other hand, that it facilitates reward-directed behaviour, making actions more precise (Bogacz, 2020; Friston et al., 2012). According to the first view, dopamine is a Pavlovian signal driving movement when reward expectation is high (Beierholm et al., 2013). In the second view, dopamine, by signalling the future reward on offer, might influence the efficiency and precision of any reward-guided behaviours based on the potential benefit accrued from rapidly and successfully obtaining that reward (Hamid et al., 2016; Manohar et al., 2015). In addition, there is evidence that cue-driven changes in dopamine levels are themselves shaped by action initiation (Coddington & Dudman, 2018; Hughes et al., 2020; Phillips, Stuber, Helen, Wightman, & Carelli, 2003; Roitman, Stuber, Phillips, Wightman, & Carelli, 2004; Syed et al., 2016). Therefore, it is also possible that the balance of dopamine transmission might instead be critical to regulate when to transition to reward-seeking, particularly when actions are not simply directed at reward (“distal” actions) or stereotyped (Nicola, 2010; Robbins & Everitt, 2007; Walton & Bouret, 2019). When we are not engaged in reward-seeking behaviour, behavioural control is reduced, making actions more variable and therefore less precise (Costa, Mitz, & Averbeck, 2019; Humphries, Khamassi, & Gurney, 2012). We term this controlled engagement for reward, behavioural “focus”. Behavioural focus in the form of cognitive control may also be governed by dopamine (Fallon et al., 2015; Westbrook & Frank, 2018).

One method to adjudicate between these accounts is to examine whether manipulating dopamine transmission differentially affects behavioural control in a context where cues on some occasions signal a requirement to make a response to gain reward and on other occasions to withhold responding to gain reward. We therefore trained rats on a symmetrically rewarded Go/No-Go task and investigated the effects of pharmacological stimulation and blockade of D1Rs, first systemically and then locally in the NAcC. By including an equal number of Go and No-Go trials associated with either high or low reward sizes, we could compare situations where a trial was either better or worse than the average reward expectation.

Across all experiments we found that Go, but not No-Go trial accuracy, was improved when the large reward was on offer. Stimulation of D1Rs consistently and rapidly biased animals to initiate actions following cue presentation, both on Go trials when responding was appropriate and crucially also on No-Go trials when responding should have been withheld. Similarly, D1R blockade disrupted response execution on Go trials (but with no influence on No-Go performance). However, while this manifested as a general reduction in the vigour of the initial actions in a sequence when the D1R antagonist was administered systemically, this was not consistently observed when it was infused directly into the NAcC. Instead, intra-NAcC D1R blockade selectively increased the *likelihood of failure* to respond appropriately on Go trials even though vigour when correctly completing a Go trial was unchanged. The response patterns on failed Go trials closely mirrored response failures off drug. Together this suggests that NAcC D1Rs normally play a key role in enabling reward expectations to regulate and focus reward seeking actions.

## Results

To allow us to investigate how dopamine transmission mediates the influence of reward over behavioural control, rats were trained on an operant Go/No-Go task which required them either to make (Go) or withhold (No-Go) action in order to gain either a small or large reward (Syed et al., 2016; Fig. 1, 2a,b). Trials were initiated by the animal entering the nosepoke, which after a short delay resulted in one of 4 auditory cues to be presented. The identity of the cue instructed them either to leave the nosepoke and respond on the left or right lever, each of which was associated with either small or large reward (side fixed for each animals, counterbalanced across animals) (***Go Small*** or ***Go Large***) or to remain in the nosepoke for the holding period in order to gain either a small or large reward (***No-Go Small*** or ***No-Go Large***). Correct performance (selecting the cued lever and pressing it twice on *Go* trials or remaining in the nosepoke for the holding period on *No-Go* trials) resulted, after a 1s delay, in delivery of reward to a magazine on the opposite wall of the operant chamber.

**Figure. 1.**
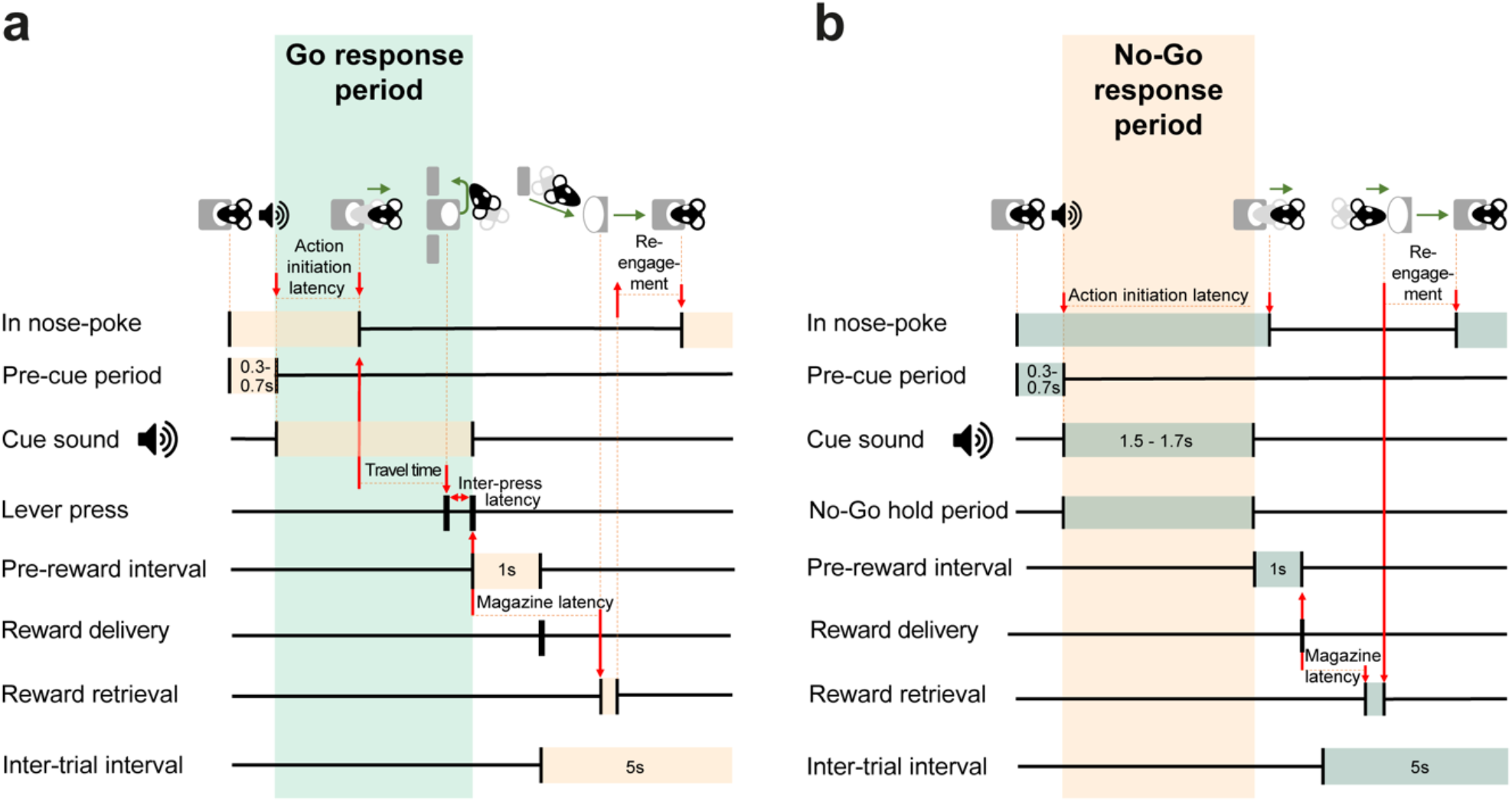
Schematic illustrating the sequence of events and associated metrics in correctly executed Go and No-Go trials. **(a)** Sequence of events and measured latencies in Go trials. Orange shading indicates recorded latencies. Green shading indicates Go trial response period, from leaving the nosepoke to completing two lever presses successfully. **(b)** Same as in (a) but for No-Go trials. Here, green shading indicates latencies and orange shading indicates the response period, in which mice were required to stay in the nosepoke.

**Figure. 2.**
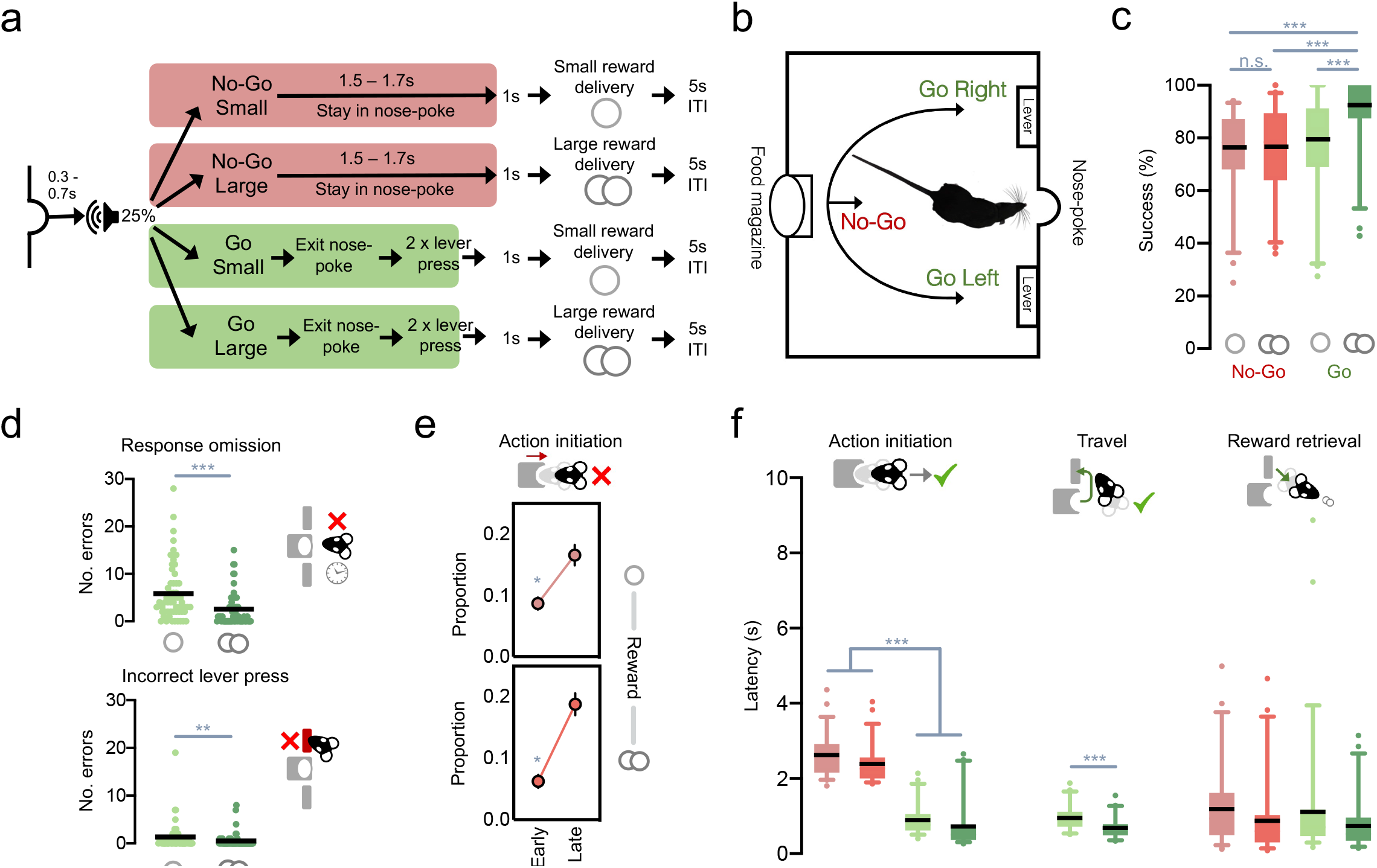
Go/No-Go task and baseline performance. **(a)** Schematic of the task trial types. Coloured shading indicates when auditory cues remained on. **(b)** Schematic of the operant chamber layout. **(c)** Animals’ performance in vehicle sessions by session split by trial type (red: No-Go; green: Go; lighter shades denote small reward trials and darker shades denote large reward trials). Solid lines indicate the mean, box extends from 25^th^ to 75^th^ percentiles, whiskers indicate 5^th^ and 95^th^ percentiles. Pairwise comparisons: Go Large vs. Go Small/No-Go Small/No-Go Large: all *p* < .001; all other comparisons n.s., *p* > .4. **(d)** Top: Total response omission errors per session. Pairwise comparison: Go Small vs. Go Large: *p* < .001. Bottom: Total incorrect lever press errors per session. Pairwise comparison: Go Small vs. Go Large: *p* = .006. **(e)** Mean proportion of times spent in the nosepoke across error No-Go trials in which animals exited early (<800ms) or late (>800ms) when a small (upper) or large (lower) reward was on offer. Pairwise comparisons: Early Small vs. Early Large: *p* = .008; Late Small vs. Late Large: n.s., *p* = .134. **(f)** Mean latencies to complete key task events in correct trials split by trial type. *** *p* < .001, ** *p* < .01, * *p* < .05

### Reward size and action requirements shape baseline performance on the task

As the current study focused more closely on behavioural measures, several of which are distinct to those reported in Syed et al. (2016), we first sought to characterise the typical performance of animals on the Go/No-Go task (Fig. 2a, b) and to determine how reliable this was across the two cohorts of rats used in the study. Pooling data across all vehicle sessions from the systemic and local experiments where all doses of the drug were administered showed that animals were able to perform well in the task (Fig. 2c), on average achieving >75% success rate across all trial types. Reward size selectively influenced response accuracy on Go but not No-Go trials (action x reward interaction: *F*_(1,56)_ = 19.455, *p* < .001). The main error type on Go trials was response omissions rather than wrong lever presses (main effect of error type: *F*_(1,56)_ = 35.183, *p* < .001), though the occurrence of both error types was decreased when the large reward was on offer (Fig. 2d; main effect of reward: *F*_(1,56)_ = 25.374, *p* < .001; error type x reward interaction: *F*_(1,56)_ = 7.834, *p* = .007).

When animals made premature responses on No-Go trials, these seldom occurred proximal to cue onset in the first, ‘early’ half of the holding period and were instead more likely in the second, ‘late’ half (Fig. 2e; main effect of No-Go period: *F*_(1,56)_ = 43.806, *p* < .001). Further, although reward size did not change the overall number of No-Go errors, it did influence when these were likely to occur, with the prospect of large reward significantly decreasing impulsive responses in the early part of the holding period but not in the late part (period x reward interaction: *F*_(1,56)_ = 6.040, *p* = .017). Reward size also had a prominent influence on response latencies on Go and No-Go trials (Fig. 2f). Behaviour was faster when a large reward was on offer, both in terms of action initiation (main effect of reward: *F*_(1,56)_ = 38.710, *p* < .001) and travel time (on Go trials: main effect of reward: *F*_(1,56)_ = 24.349, *p* < .001), as well as reward retrieval (main effect of reward: *F*_(1,55)_ = 21.171, *p* < .001; one animal excluded due to faulty magazine detector).

Importantly, although the animals in the cannulated cohort had on average slightly lower success rates on all trial types (main effect of cohort: *F*_(1,56)_ = 6.102, *p* = .017), on almost all other task latency measures there was no reliable difference across cohorts (all main effects or interactions with cohort: *F* < 2.6, *p* > .1; except for cohort x reward for the time in nosepoke metric *F*_(1,56)_ = 4.699, *p* = .034, though even here post-hoc tests showed no difference between cohorts, both *p* > .2). Taken together, this demonstrates that baseline behaviour on the Go/No-Go task across both cohorts is strongly and consistently mediated by both action requirements and reward size.

### Global D1Rs regulate action initiation and the vigour of actions distal to reward

We next investigated what role D1Rs play in modulating appropriate action restraint and action initiation for future reward by analysing the effects of systemic administration of either a D1 agonist, SKF-81297 (Cohort 1) or a D1 antagonist, SCH-23390 (Cohort 2, *see methods*).

#### No-Go trials

Systemic administration of a D1R agonist SKF-81297 had no influence on rates of aborted trials during the pre-cue hold period (main effect of drug: *F* < .5, *p* > .6, data not shown). However, it substantially impaired performance in No-Go trials (Fig. 3a; main effect of drug: *F*_(2,20)_ = 14.911, *p* < .001; drug x reward interaction: *F*_(2, 20)_ = 3.467, *p* = .051), elevating the overall number of errors (main effect of drug: *F*_(2,20)_ = 12.165, *p* < .001). As can be observed in Fig. 3c, the drug did not cause a uniform increase in the probability of making a premature response; instead, D1R stimulation selectively increased inappropriate action initiation only in the early half of the No-Go hold period (Fig. 3d; drug x error period interaction: *F*_(2,20)_ = 7.780, *p* = .003. This was effectively the opposite of the effect of reward, which reduced early leaving; no drug x reward x period interaction, *p* > .3). On correct No-Go trials, when animals had successfully withheld responding during the No-Go period, D1R stimulation did not change the overall speed of initiation but did reduce the difference between latencies in small and large reward trials (Fig. 3b; drug x reward interaction: *F*_(2,20)_ = 7.264, *p* = .004; main effect of drug n.s., *F* < .5, *p* > .6) although there was no corresponding effect on reward collection (*F* < 1.9, *p* > .1, data not shown).

**Fig. 3.**
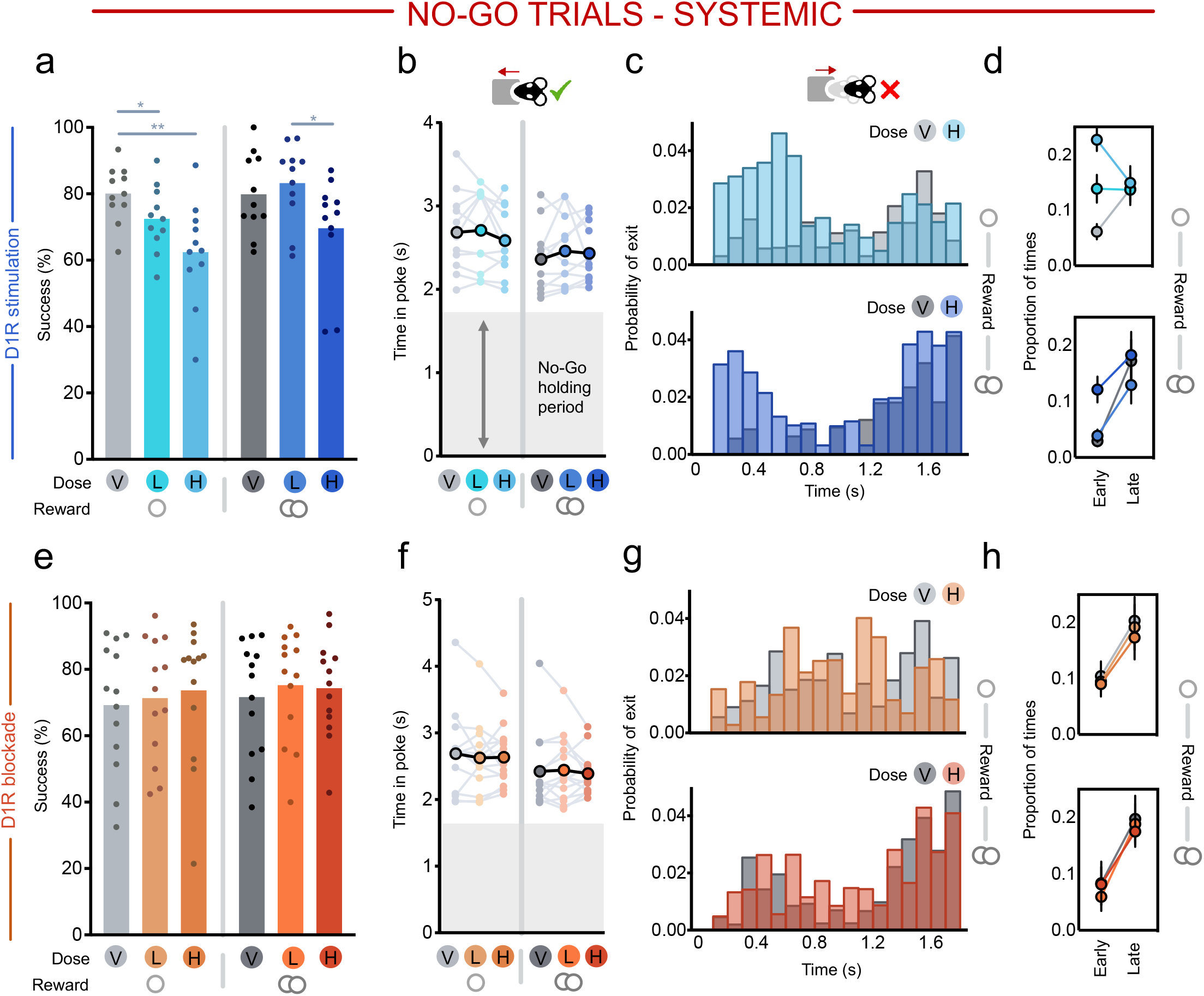
Systemic effects of D1R stimulation (SKF-81297) or blockade (SCH-23390) in No-Go trials. V = vehicle, L = low dose, H = high dose. Single circle indicates small reward condition, double circle indicates large reward condition. **(a-b)** Effects of D1R stimulation split by small (left) and large (right) reward No-Go trials on **(a)** success rate and **(b)** time in nosepoke in successful trials. For b, analysis of pairwise comparisons due to significant drug x reward interaction: vehicle small reward vs. large reward: *p* = .005, low dose small reward vs. large reward: *p* = .012, high dose small reward vs. large reward: *p* = .071. **(c)** Mean probability histogram of time in nosepoke in failed small (upper) and large (lower) reward No-Go trials for saline (grey) or high dose (blue) manipulations, calculated as probability over all head exit times. **(d)** Mean proportion of times spent in the nosepoke across trials in which animals exited early (<800ms) or late (>800ms) when a small (upper) or large (lower) reward was on offer. Pairwise comparisons: early period vehicle vs. low dose: *p* = .003, vehicle vs. high dose: *p* < .001; late period, all *p* > .5. **(e-h)** Same as in **(a-d)** but for systemic D1R blockade. \*\**p* < .01, \**p* < .05

Taken together, these results demonstrate a role of activity at global D1Rs in promoting early cue-driven action both when a small or a large reward was on offer. However, this effect was asymmetric as systemic D1R blockade with the antagonist SCH-23390 had no significant effect on No-Go performance (Fig. 3e; no main effect of drug, reward, or interaction: all *F* < .6, *p* > .4), latencies to leave the nosepoke (Fig. 3f-h; all *F* < .7, *p* > .5) or time taken to collect reward in successful trials (all *F* < 1.0, *p* > .4).

#### Go trials

We next sought to examine the influence of D1Rs on the accuracy and speed of action on Go trials. Unexpectedly, both systemic D1R stimulation *and* D1R blockade impaired performance on Go trials.

The D1R agonist reduced success rate selectively on Go Small trials at the highest dose (Fig. 4a; drug x reward: *F*_(2,20)_ = 4.135, *p* = .031). This was caused both not only by a numeric increase in response omissions on Go Small trials (Fig. 4b; drug x reward: *F*_(2,20)_ = 3.346, *p* = .056), but also by a small but reliable increase in the number of wrong lever errors on Go Small trials (i.e., high reward lever) (Fig. 4c; drug x reward: *F*_(2,20)_ = 4.515, *p* = .024). Although the D1R agonist numerically speeded animals’ latency to exit the start poke on small reward trials (Fig. 4d; *F*_(2,20)_ = 2.775, *p* = .086), it *slowed* travel time from head exit to a correct lever response (Fig. 4e; main effect of drug: *F*_(2,20)_ = 6.331, *p* = .007), in line with greater response competition from the high reward lever. Subsequent trial re-initiation latencies after success were also slower (main effect of drug: *F*_(2,20)_ = 11.954, *p* < .001).

**Figure 4.**
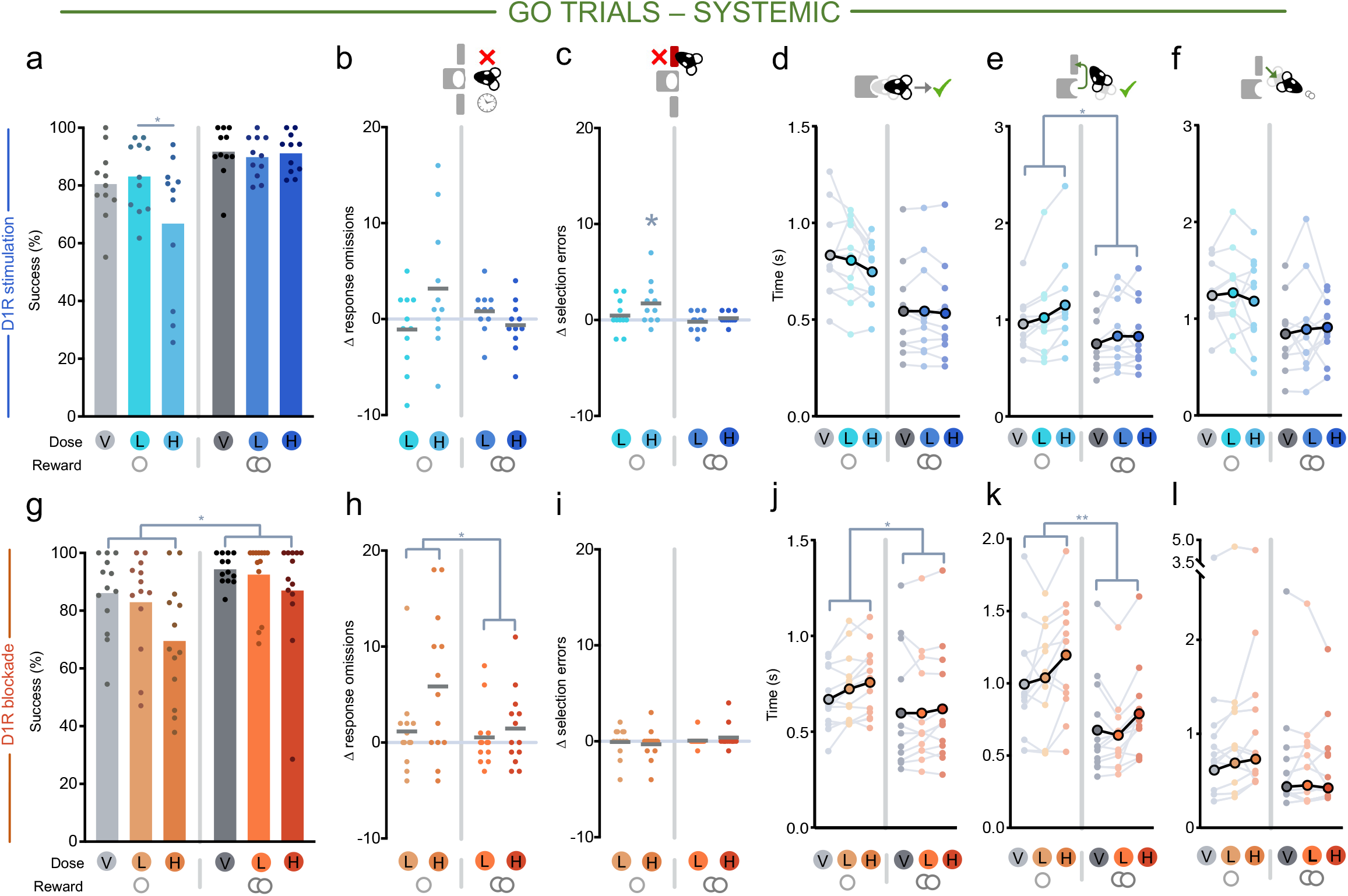
Systemic effects of D1R stimulation (SKF-81297) or blockade (SCH-23390) in Go trials. V = vehicle, L = low dose, H = high dose. Single circle indicates small reward condition, double circle indicates large reward condition. **(a-f)** Effects of local D1R stimulation split by small (left) and large (right) reward Go trials on **(a)** success rate, **(b)** response omission errors (relative to vehicle session), **(c)** lever selection errors (relative to vehicle session), **(d)** latency to leave the nosepoke after Go cue onset, **(e)** latency from nosepoke exit to first lever press, **(f)** and latency from trial completion to entering the food magazine to retrieve reward. **(g-l)** Same as in **(a-f)** but for systemic D1R blockade. \*\**p* < .01, \**p* < .05

The D1R antagonist also caused a dose-dependent reduction in Go trial success rate (Fig. 4g; main effect of drug: *F*_(2,24)_ = 7.015, *p* = .004; drug x reward interaction n.s, *F* < 1.1, *p* > .3). However, this was driven primarily by increased response omissions (Fig. 4h; main effect of drug: *F*_(2,24)_ = 6.846, *p* = .004; drug x reward interaction, *F* < 2.9, *p* > .07) and there was no effect on the ability to select the correct lever (Fig. 4i; no main effect or interaction of drug: *F* < .9, *p* > .4). D1R blockade also slowed latencies, but this was evident for *all* distal elements – i.e. all actions aside from direct approach to the food magazine – of the Go trial sequence: exiting the start poke (Fig. 4j; main effect of drug: *F*_(2,24)_ = 8.607, *p* = .002; drug x reward interaction: *F*_(2,24)_ = 2.903, *p* = .074), travelling to the lever (Fig. 4k; main effect of drug: *F*_(2,24)_ = 13.226, *p* < .001), and reinitiating the subsequent trial after success (main effect of drug: *F*_(2,24)_ = 12.231, *p* < .001, data not shown). However, there is indication that this was not a non-specific motoric effect as the drug did not significantly slow time to retrieve reward following successful trial completion (Fig. 4l; no main effect or interaction with drug: both *F* < 2.1, *p* > .15).

In sum, as with No-Go trials, we again find an asymmetric effect of stimulation and blockade of D1Rs. But here, whilst the D1R agonist affected animals’ ability to efficiently perform the correct action – as also demonstrated by the increase in wrong lever responses and slower travel times – the D1R antagonist more broadly slowed actions outside of directly travelling to retrieve reward such that animals increasingly omitted responding. This influence of D1Rs on rapid cued action and the vigour of actions distal to reward appeared specific to this receptor, as systemic administration of a D2R agonist instead slowed all Go and No-Go latencies (Supp. Text 1, Supp. Fig. 1).

### D1Rs in NAcC selectively shape action likelihood and focus

The first experiments demonstrated a key selective role for D1Rs in rapid modulation of action restraint and initiation. As our previous study had demonstrated a close relationship between fast increases in dopamine levels in NAcC and action initiation (Syed et al., 2016), our overall hypothesis was that D1Rs in NAcC would be a critical locus for this. In particular, we hypothesised that on No-Go trials, stimulating D1Rs in the NAcC would promote action over inaction, causing an increase in fast premature errors on No-Go trials and reducing latencies to initiate responding on Go trials. By contrast, antagonism of this receptor subtype would have little effect on No-Go trials (as endogenous dopamine is already suppressed on these trials), but would slow responding on Go trials. Furthermore, based on previous work showing that mesolimbic dopamine has a limited role in selecting *between* actions, particularly when the required response paths are fixed (Hollon, Arnold, Gan, Walton, & Phillips, 2014; Nicola, 2010), we reasoned there should be no change in the type of errors made or how animals executed actions in Go trials. Therefore, we examined the effects of infusions of either the D1R agonist or antagonist directly into the NAcC (cohort 2). To ensure consistency with the effects we observed in the first cohort, prior to surgery we replicated the systemic D1R agonist experiment and found a comparable pattern of effects on No-Go and Go performance (Supp. Fig. 2; drug x cohort interactions: all *p* > .2).

#### No-Go trials

On No-Go trials, intra-NacC administration of a D1R agonist or antagonist replicated the majority of the effects of systemic administration. Specifically, NAcC D1R stimulation increased premature responses after cue onset on No-Go trials (Fig. 5a; main effect of drug: *F*_(2,24)_ = 8.459, *p* = .002) and this was again particularly evident early in the No-Go holding period, although here the highest dose also increased errors in the late period (Fig. 5c, d; main effect of drug: *F*_(2,22)_ = 6.630, *p* = .006; drug x period interaction: *F*_(2,22)_ = 3.613, *p* = .044). On correctly performed No-Go trials, as before, there were no reliable changes in the speed to exit the nosepoke (Fig. 5b) or to reach the magazine (all *F* < 2.7, *p* > .09).

**Fig. 5.**
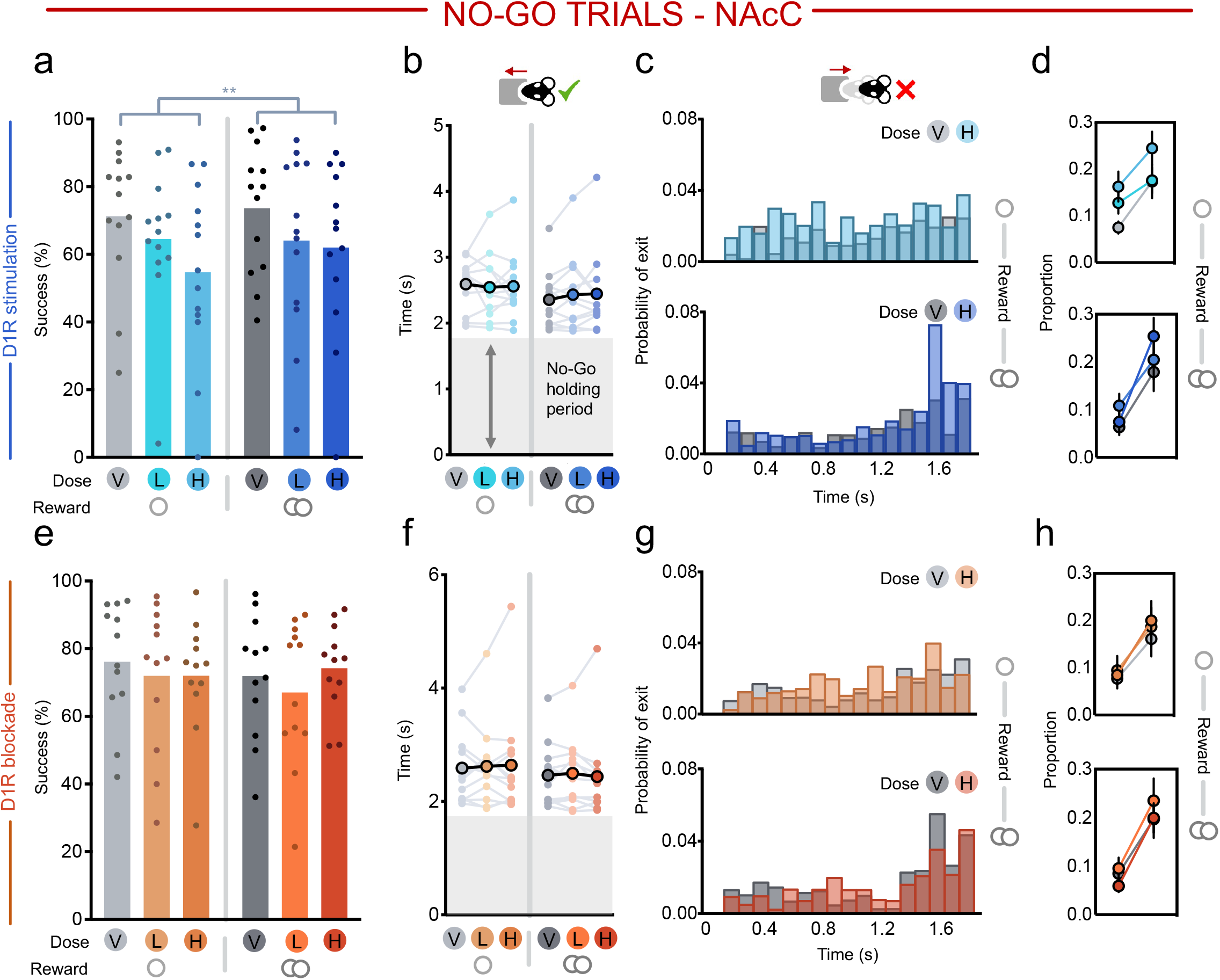
Effects of intra-NAcC D1R stimulation (SKF-81297) or blockade (SCH-23390) in No-Go trials. V = vehicle, L = low dose, H = high dose. Single circle indicates small reward condition, double circle indicates large reward condition. **(a-b)** Effects of D1R stimulation split by small (left) and large (right) reward No-Go trials on **(a)** success rate and **(b)** and time in nosepoke in successful trials. **(c)** Mean probability histogram of time in nosepoke in failed small (upper) and large (lower) reward No-Go trials for saline (grey) or high dose (orange and red) manipulations, calculated as probability over all head exit times. (Pairwise comparisons: early period vehicle vs. low dose: *p* = .019, vehicle vs. high dose: *p* = .022; late period vehicle vs. high dose: *p* = .009, vehicle vs. low dose: n.s., *p* > .6). For this analysis we excluded 1 animal where on average > 50% of the errors occurred in the early No-Go period, which was > 3 S.D. from the group. **(d)** Mean proportion of times spent in the nosepoke across trials that were early (< 800ms) or late (> 800ms) for small (upper) and large (lower) reward trials. **(e-h)** Same as in **(a-d)** but for local D1R blockade. \*\**p* < .01, \**p* < .05

To investigate what was causing this increase in premature errors, we used video tracking on a subset for rats for which we were able to perform video analyses (n = 6, see *Methods*) to establish the behaviour of the rats in these erroneous No-Go trials (Supp Fig 3a, b). This revealed that rats were more likely to directly visit the food magazine than either lever, particularly when a large reward was on offer (Supp. Fig. 3c-e; main effect of *F*_(2,8)_ = 13.448, *p* = .003; location x reward interaction: *F*_(2,8)_ = 4.899, *p* = .041). Importantly, this response pattern was comparable after intra-NAcC D1R agonist administration (Supp. Fig. 3c-e; main effect of drug, drug x reward x location interaction, both *F* < 1.6, *p* > .25), the only difference being that the drug tended to reduce the likelihood of reaching any target location on small reward trials (drug x reward interaction: *F*_(1,4)_ = 27.495, *p* = .006). Therefore, although stimulation of NAcC D1Rs increased the *likelihood* of premature No-Go responses, this was not driven by a selective change in responses towards the levers or food magazine.

By contrast, intra-NAcC infusion of the D1R antagonist had no effect on performance or latencies in No-Go trials, replicating the pattern of results from systemic administration (Fig. 5e-h; all *F* < 1.6, *p* > .2). This implies that NAcC D1R stimulation rapidly promotes action over inaction in the presence of reward-associated cues, even though here this is disadvantageous.

#### Go trials

The effect of intra-NAcC administration of the D1R agonist or antagonist had more selective effects on Go trials than was observed after systemic administration. Stimulation of NAcC D1Rs, unlike systemic administration, had no overall effect on the proportion of correct responses on Go trials (Fig. 6a; main effect of drug and interaction: both *F* < 2.3, *p* > .1). It did, however, promote faster action initiation (Fig. 6d; main effect of drug: *F*_(2, 24)_ = 4.046, *p* = .031), although, unlike with systemic administration, neither the speed with which animals travelled to the lever or retrieved the reward were affected (Fig. 6e, f; both *F* < .9, *p* > .4). This further supports a role for NAcC D1R stimulation in the rapid promotion of action initiation as only the speed to initiate the action sequence was altered.

**Fig. 6.**
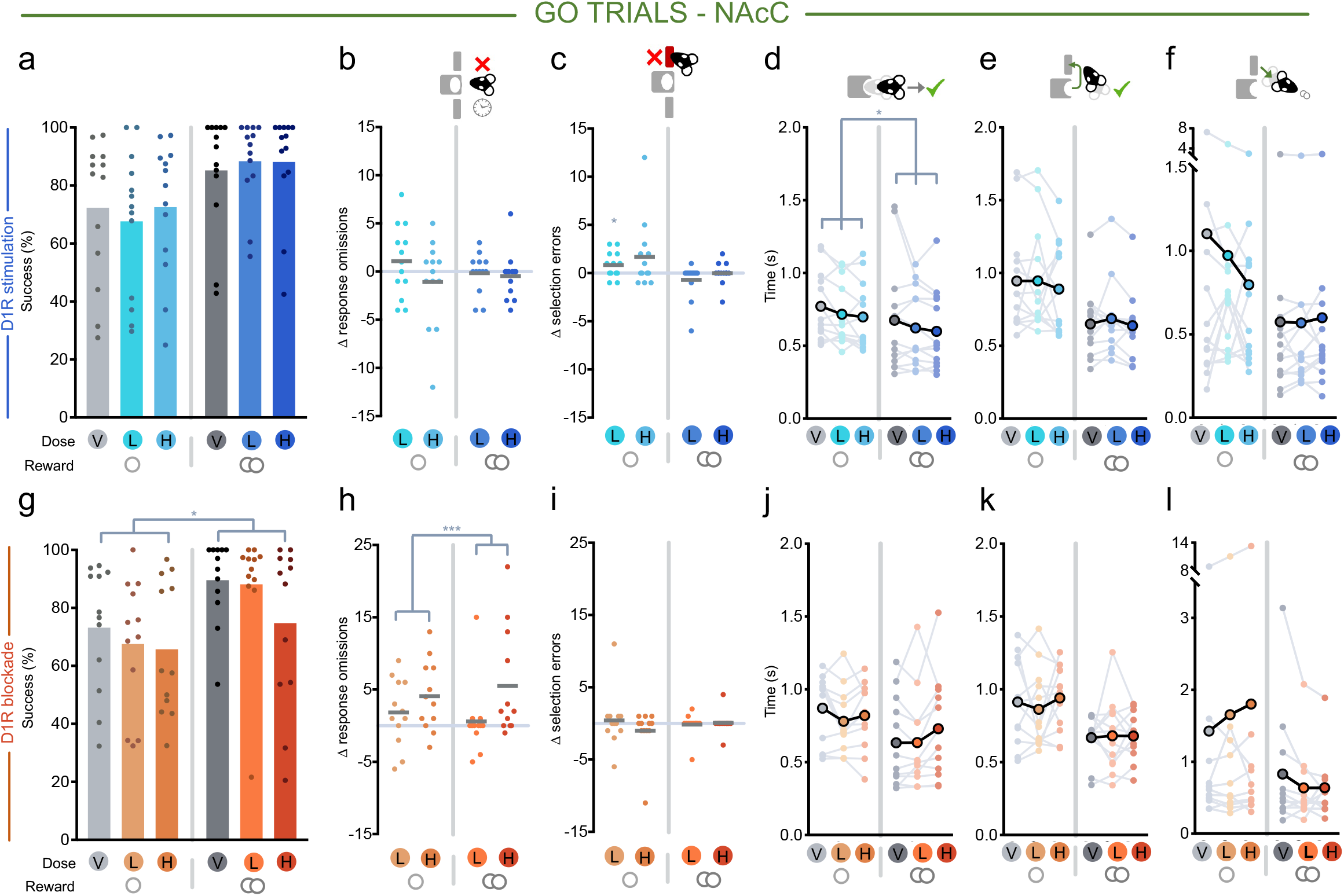
Effects of intra-NAcC D1R stimulation (SKF-81297) or blockade (SCH-23390) in Go trials. V = vehicle, L = low dose, H = high dose. Single circle indicates small reward condition, double circle indicates large reward condition. **(a-f)** Effects of local D1R stimulation split by small (left) and large (right) reward Go trials on **(a)** success rate, **(b)** response omission errors, **(c)** lever selection errors, **(d)** latency to leave the nosepoke after Go cue onset, **(e)** latency from nosepoke exit to first lever press, **(f)** and latency from trial completion to entering the food magazine to retrieve reward. **(g-l)** Same as in **(a-f)** but for local D1R blockade. \*\**p* < .01, \**p* < .05

Blockade of NAcC D1Rs resulted in a lower success rate in Go trials, mirroring the effect with systemic administration (Fig. 6g; main effect of drug: *F*_(2, 22)_ = 4.559, *p* = .022), and this was again caused by a selective increase in response omissions (Fig. 6h; main effect of drug: *F*_(2,22)_ = 4.542, *p* = .022; lever selection errors both *F* < 1.9, *p* > .18; Fig. 6i). However, whereas systemic D1R blockade had significantly slowed distal latencies, here, surprisingly, intra-NAcC administration of the D1R antagonist did not affect any latencies – action initiation, travel time, and reward collection (Fig. 6j-l; no main effects or interactions with drug, all *p* > .09).

#### Focused responding on Go trials is shaped by reward and is mediated by NAcC D1Rs

These data demonstrate a conspicuous and surprising dissociation of intra-NAcC D1R blockade between the disruption of successful Go trial completion within a 5s time window (Fig. 6g-h) coupled with an absence of effect on the speed of responding on correctly performed Go trials (Fig 6j-l). To understand this better, we examined in more detail the pattern and performance on Go trials on and off intra-NAcC D1R blockade. First, we investigated whether this dissociation could be caused by the intra-NAcC D1R antagonist having a cumulative effect on arousal within a session. We reasoned that if this was the case, on drug, the correct responses with normal response latencies may predominate at the beginning of the session and the response omissions may cluster later in the session. In fact, however, these elevated error rates were equally distributed across the session in both vehicle and drug sessions and a difference in response trial omission rates was already apparent in the first quartile of the session (Fig. 7a; main effect of drug: *F*_(2,22)_ = 4.609, *p* = .021; no main effect of quartile or interaction, both *F* < .7, *p* > .5).

**Fig. 7.**
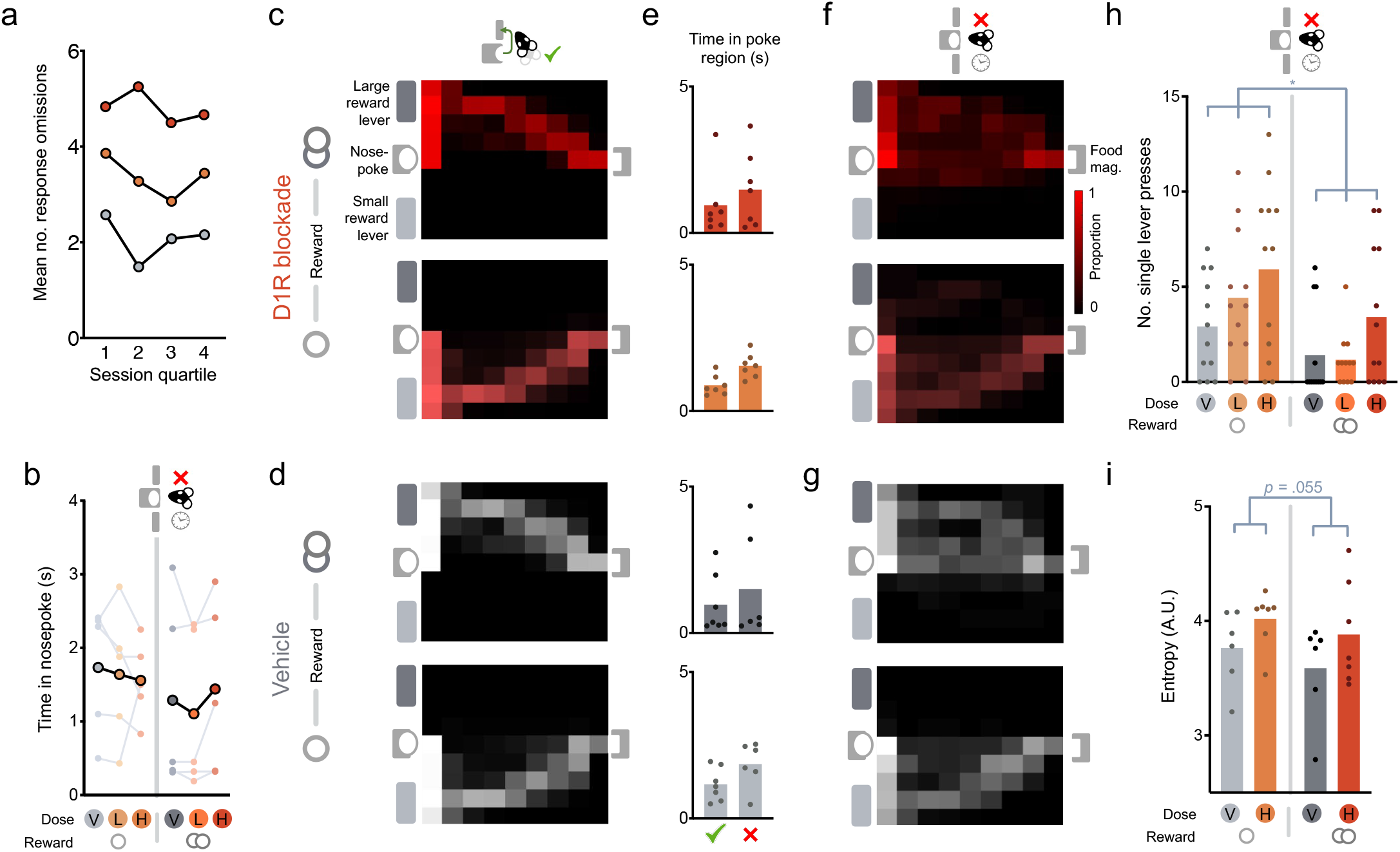
The effects of intra-NAcC D1R blockade (SCH-23390) in response omission Go trials. **(a)** Mean number of response omission errors made across rats across sessions when each session is split into quartiles. **(b)** Mean time in nosepoke from cue onset in response omission trials. **(e)** Mean time in the area of the nosepoke (see *Methods: Video Analyses*). **(c, d, f, g)** Mean probability density across rats in small (lower) or large (upper) reward Go trials, when **(c)** correct on the high dose of the intra-NAcC D1antagonist, **(d)** correct on vehicle, **(f)** in response omission trials on the high dose of the intra-NAcC D1 antagonist and **(g)** in response omission trials on vehicle. **(h)** Total number of single lever presses in response omission trials. **(i)** Average entropy of animals in response omission trials. Data displayed for all animals for which we had tracking, but statistical analysis was restricted to n=5 for which we had a reliable tracking in both drug and vehicle sessions.

Next, we examined whether the drug caused rats’ responding on these omission trials to be more likely to be disordered. We reasoned that this could manifest in three ways: (1): “opting out”, staying near the start port and waiting for the next trial; (2) “incorrect cue detection”, revealed by an increase in trajectories to the wrong lever; or (3) “unfocused”, where the appropriate action is taken, but with less vigour and accuracy, thereby resulting in the rat failing to meet the response requirement of the trial. To assess this, we again examined response variables and within-trial trajectories using video tracking on a subset of rats (n=7, see *Methods*; note that for analyses comparing within-subject changes in performance on and off drug for both reward sizes, n=5 due to 2 animals not making response omission errors in the saline condition. Replicated analyses when averaged across rewards to give n = 7 result in the same direction of effects in all cases), focusing on comparisons between intra-NAcC administration of the high dose of the D1R antagonist or vehicle.

While animals were overall slower to initiate actions on omission trials in comparison to correctly performed Go trials, importantly this was no different with or without intra-NAcC D1R blockade (Fig. 7b; main effect of outcome: *F*_(1,4)_ = 11.816, *p* = .026; no main effect of drug or interaction with outcome or reward, all *F* < 1.5, *p* > .2; if only small reward trials analysed to account for the low error rates on high reward trials on vehicle, main effect of outcome: *F*_(1,9)_ = 13.328, *p* = .005; no main effect of drug or interaction with outcome, all *F* < .9, *p* > .4). Similarly, time spent in a defined area near the nosepoke after erroneous head exits in Go trials was unchanged by the intra-NAcC D1R antagonist, suggesting that rats were not “opting out” (Fig.7e; no main effect of drug or interaction, both *F* < 1.0, *p* > .3).

In fact, during the 5s cue presentation on these omission trials, rats not only moved away from the nosepoke, but they would often perform similar sequences of actions as on correct Go trials – moving towards the cued lever and even subsequently heading to the food magazine (Fig. 7c-f). This suggests that rats were not suffering from erroneous cue detection. Strikingly this pattern was equivalent whether or not they had been administered the D1R antagonist or vehicle, despite the fact that the overall propensity of rats to make omission errors was increased with the antagonist. Specifically, the proportion of omission trials in which rats first visited the region of the correct lever was significantly higher in comparison to first visiting the incorrect lever, but this was unaltered by the drug (main effect of outcome: *F*_(1,4)_ = 100.791, *p* = .001; no main effect of drug, reward, or interactions, all *F* < .5, *p* > .4; average proportion of correct lever responses: vehicle small reward: 0.72 ± 0.11, large reward: 0.75 ± 0.14; SCH small reward: 0.65 ± 0.09, large reward: 0.65 ± 0.15, mean±SEM) and the cumulative probability of visiting the area near the correct lever when on drug did not significantly differ from vehicle (no main effect of drug or interaction, both *F* < .4, *p* > .5). There was also no difference due to drug in how likely the rats were to visit the correct lever and then go on complete the trajectory by visiting the magazine (no main effect or interaction with drug, both *F* < .5, *p* > .5; vehicle small reward: 0.42±0.11, large reward: 0.53±0.21; SCH small reward: 0.39±0.14, large reward: 0.42±0.15). Overall, trajectory lengths during the 5s cue window were comparable between error and correct trials on or off drug (no main effect of drug or interaction, both *F* < 1.2, *p* > .3).

Yet importantly, although the trajectories on omission trials contained many features common with correctly performed Go trials, responding on omissions nonetheless lacked equivalent focus and precision.

This is in part demonstrated by the fact that in omission trials they were more likely to make a single response on the correct lever rather than the two required for the trial to be successful (Fig. 7h; main effect of drug: *F*_(2,22)_ = 5.571, *p* = .011). Moreover, the entropy, or noisiness, of the animals’ trajectories in omission trials on and off drug showed a strong trend for entropy to be increased by the intra-NAcC D1R antagonist (Fig. 7i; main effect of drug: *F*_(1,4)_ = 7.201, *p* = .055).

Together this suggests that the promise of reward, signalled by cues, facilitates animals to engage in focused reward-seeking sequences through NAcC D1Rs and that blockade of these signals reduces the likelihood of animals transitioning to this focused reward-seeking state.

## Discussion

Dopamine transmission is a key component mediating the influence of reward predictions on behaviour, yet its precise role in cue-driven behavioural control has remained contentious (Averbeck & Costa, 2017; Gershman & Uchida, 2019; Robbins & Everitt, 2007; Salamone & Correa, 2012; Walton & Bouret, 2019). Here we used a factorial design, which separately manipulated the size of the reward on offer and the behavioural requirements to gain that reward, to investigate the role of dopamine transmission at D1Rs in regulating this relationship. Stimulation, but not blockade, of D1Rs across the whole brain or locally in the NAcC consistently disrupted No-Go performance, potentiating inappropriate responses that clustered strongly just after cue presentation. The most prominent effect of D1R blockade, by contrast, was to increase response omissions on Go trials. While this manifested as a general reduction of invigoration of all distal actions in the response sequence after systemic administration (action initiation and travel time latencies, but not reward collection), this was not observed after intra-NAcC blockade where on correctly performed trials these metrics were unaffected. Instead, the disruption of transmission at NAcC D1Rs increased the probability that Go trial performance was in an “unfocused” state, characterised, both on and off drug, by a reduction in the precision of responding even though the appropriate action sequence was often executed.

The prospect of reward can positively shape both the speed and precision of behaviour (Guitart-Masip, Duzel, Dolan, & Dayan, 2014; Kawagoe, 1998; Manohar et al., 2015; Shadmehr, Reppert, Summerside, Yoon, & Ahmed, 2019), and several lines of evidence suggest that dopamine may play a key role in mediating aspects of both processes (Beierholm et al., 2013; Hamid et al., 2015; Manohar et al., 2015; Niv et al., 2007; Westbrook et al., 2020). As expected, rats’ performance in the current experiment was also strongly affected by the reward size on offer. Cues associated with a large future reward reduced latencies to initiate actions and to complete each prerequisite element of the action sequence (the correct lever on a Go trial and, on both trial types, the food magazine).

This finding is consistent with the notion that there is a direct link between the vigour of actions – the reciprocal of the time to complete an action sequence (Shadmehr et al., 2019) – and the net gain from obtaining the potential reward (Niv et al., 2007; Pompilio & Kacelnik, 2010; Shadmehr, Huang, & Ahmed, 2016). However, there was an asymmetric influence on response accuracy, with the prospect of a large reward improving Go trial accuracy by reducing the likelihood that animals would fail to make a response in the allotted time window, but having no reliable effect successful No-Go trial completion. This could be caused by reward having distinct influences on separable processes during No-Go trials, boosting not only instrumental precision but also a Pavlovian influence to approach rewarded locations, which here is maladaptive (Lex & Hauber, 2010). Indeed, in No-Go trials, where animals exited the nosepoke prematurely we found that the rats were more likely to approach the food magazine, particularly when the large reward was on offer (Supplementary Fig. 3). A related mechanistic alternative is that the rats have learned through action to limit how reward modulates cue-driven dopamine on No-Go trials to avoid premature responses. While the presentation of cues associated with future reward can rapidly increase dopamine levels in terminal regions in relation to the value of available reward (or, more specifically, the *change* in benefit signalled by the cue compared to previous expectation) (Gan et al., 2010; Tsutsui-kimura et al., 2020), we and others have found using fast-scan cyclic voltammetry that release patterns are suppressed until a reward-seeking action is made to gain that benefit (Roitman et al., 2004; Syed et al., 2016).

What is in no doubt though is that pharmacological stimulation of D1Rs rapidly promoted actions to be initiated, typically speeding action initiation on Go trials, but also consistently increasing inappropriate No-Go responses. Notably, these latter premature actions were most evident early in the No-Go holding period just after cue presentation; but if the animal was able to withhold responding at this point, it was often no more likely to make an error in the second half of the holding period than off drug. Moreover, neither systemic nor intra-NAcC D1R stimulation caused an increase in head exits during the pre-cue period, implying that it was cue presentation that elicited the behavioural response.

While these findings are generally consistent with studies implicating hyperdopaminergic states with an increased likelihood of motor or ‘waiting’ impulsivity (Pattij et al., 2007; Pezze et al., 2007), it is important to note that the mechanisms of behavioural control taxed in the current task, where animals have to suppress responding before and during the presentation of a reward-associated cue, may well be distinct from those in tasks such as the 5-choice serial reaction time task (5-CSRTT), which requires animals to wait until a cue is presented. For example, D1R stimulation does not always increase premature responses in the 5-CSRTT (Passetti, Levita, & Robbins, 2003; Pezze et al., 2007). Conversely, intra-NAcC D1R blockade has been shown to reduce premature responses on the 5-CSRTT (Pattij et al., 2007), but here had no effect on No-Go performance. This demonstrates that although activity at D1Rs can promote cue-driven decisions to act, it is not necessary for actions to be executed. Finally, we have reported that intra-NAcC administration of the stimulant amphetamine causes a much broader range of premature responses than observed in the current study, with increases in impulsive actions observed not only throughout the early and late intervals of the No-Go holding period but also in the pre-cue period (Harmson, Grima, Panayi, Husain, & Walton, 2020).

Overall, our data support the idea that D1Rs, likely mediated by those within the NAcC, enable cues signalling reward opportunities to promote transitions to action. This is consistent with findings that cue-evoked excitation of D1-expressing MSNs is closely tied to the latency to initiate reward-seeking behaviour (du Hoffmann & Nicola, 2014; Nicola, 2010; Saunders, Richard, Margolis, & Janak, 2018). Of particular relevance, in one recent study, du Hoffmann and Nicola showed that intra-NAcC administration of D1 agonists promoted the likelihood of cue-driven behaviour for sucrose reward in a state of satiety (du Hoffmann & Nicola, 2016), which several groups, including our own, have shown attenuates dopamine release to cues signalling the potential availability of sucrose reward (Aitken, Greenfield, & Wassum, 2016; Papageorgiou et al., 2016). Moreover, as in these previous studies, it appears that NAcC D1Rs play a specific role in invigorating the initiation of an action sequence, but then have little influence over the vigour of ongoing actions, with the time to reach the lever or collect the reward unaffected by either D1R blockade or inhibition. This contrasts with the effects of systemic manipulation of D1Rs, which not only affected initiation latencies but also the speed of lever approach (though not reward retrieval). One possibility is that regulation of the movement vigour, particularly in the service of gaining response-contingent rewards, relies on D1Rs in dorsal striatum (Baraduc, Thobois, Gan, Broussolle, & Desmurget, 2013; Grogan, Sandhu, Hu, & Manohar, 2020; Panigrahi et al., 2015). Notably, both optogenetic inhibition and stimulation of substantia nigra pars compacta dopamine cells or D1-expressing MSNs have been shown to disrupt ongoing movements (Bova et al., 2020; Tecuapetla, Jin, Lima, & Costa, 2016), which parallels the effect observed here that systemic administration of not just the D1R antagonist but also the D1R agonist slowed travel to the lever. The latter manipulation also caused a small but reliable increase in incorrect lever presses on Go trials, and both effects may reflect competition between different potential reward-associated instrumental responses in dorsal striatum (Bova et al., 2020).

Given the importance of NAcC D1Rs in regulating decisions to act, it might initially seem entirely expected that intra-NAcC D1R blockade would also cause an increase in the proportion of response omissions on Go trials, comparable to what had been observed after systemic administration. However, two aspects of this make it more surprising. First, a number of elegant experiments have shown that NAcC dopamine transmission is only important for flexible or taxic responses – in other words, when needing to take a novel path to gain reward (Nicola, 2010) – yet here the start and goal locations are fixed across trials. Second, this increase in omissions occurred alongside an absence of an effect on any latency measures on correctly performed trials. This could not be explained by a simple arousal effect causing animals to become increasingly amotivated over time, for instance if the D1R antagonist reduced the efficacy of rewards to sustain behaviour (Fischbach & Janak, 2019) or the animal’s intrinsic motivation was reduced by satiety (du Hoffmann & Nicola, 2016; Papageorgiou et al., 2016), as omission error rates were comparable from the start to the end of the session. Moreover, there was no evidence that the rats were simply disorganised or disengaged during omissions after D1R administration; not only was there no increase in wrong lever choices but also, strikingly, analysis the patterns of responding in a subset of animals on these trials showed that they performed many of the same action sequence components observed on correctly performed Go trials including movement to the cued lever and, on a notable proportion of trials, then towards the food magazine as if to retrieve reward.

Instead, what characterised performance on response omissions was a marked reduction in vigour and precision in the execution of the response sequence: slower initiation, less focused responses towards the correct lever, increased likelihood of only making one of the two required lever presses. Crucially, this unfocused state had not emerged *de novo* with administration of the intra-NAcC D1R antagonist, but instead was a potentiation of an analogous response pattern also observed off drug. Response omissions in baseline sessions most commonly occurred on small reward trials, which generate an initial dip in NAcC dopamine (Syed et al., 2016). This suggests that endogenous rapid dopamine release, such as occurs when cues signal an improved reward opportunity, play a key role in promoting transitions to focused and efficient responding. In the absence of dopamine, it becomes more likely that animals will act in an unfocused state, as was demonstrated by the increase of single lever presses and overall entropy when animals omit responding in Go trials, which fails to ensure each element of the required sequence is completed efficiently and in order. This may be relevant for understanding the actions of therapeutic doses of stimulant drugs such as amphetamine, which can potentiate evoked NAcC dopamine and increase sustained attention (Andrzejewski et al., 2014; Schuweiler, Athens, Thompson, Vazhayil, & Garris, 2018). Nonetheless, it is important to note that stimulation of NAcC D1Rs did not concomitantly increase the success rate on Go Small reward trials. This demonstrates that whilst D1R transmission is necessary to facilitate transitions to focused reward seeking, it is not sufficient in the absence of other inputs.

Pronounced changes in dopamine can increase the excitability of D1-expressing MSNs (du Hoffmann & Nicola, 2014; Lahiri & Bevan, 2020; Lee et al., 2020) and therefore we focused our investigations on the downstream effects of cue-elicited dopamine on D1Rs. Nonetheless, several studies have also demonstrated important roles for D2Rs for sustaining responding and neural activity in NAcC D2-expressing MSNs evoked by reward-associated cues (du Hoffmann & Nicola, 2014; Lex & Hauber, 2010; Nicola, 2010). We therefore also compared at a systemic level the effects of low-to-moderate doses of a D2/D3-receptor agonist quinpirole and a D2-receptor antagonist eticlopride. While D2 blockade had no reliable effect on any measure, stimulation of D2/D3Rs strongly disrupted Go trials, slowing all movements in the sequence, including reward collection, and at the highest dose substantially increasing the proportion of response omissions. However, it was not simply that the animals’ movement was impaired as they also showed an increase in premature head exits during the pre-cue period. This could partly reflect an effect of the agent on D2 autoreceptors causing a reduction in midbrain dopamine activity and dopamine release likelihood (Bunney, Walters, Roth, & Aghajanian, 1973; Marcott, Mamaligas, & Ford, 2014; Schmitz, Schmauss, & Sulzer, 2002), although it is notable that the effects of D2R stimulation are partially distinct from D1R (and D2R) blockade. An alternative possibility is that it reflects the fact that D2-expressing MSNs encode the anticipated costs of acting or the value of alternative options (Collins & Frank, 2014; Tecuapetla et al., 2016). However, future studies, specifically targeting NAcC D2Rs would be required to disentangle these possible explanations.

Together, these results help refine ideas about the role of mesolimbic dopamine, acting via NAcC D1Rs, in reward-guided choice: it is not required to sustain appropriate selection *between* options, but instead plays a key role in the rapid translation of information about the potential net gain of a presented opportunity into a decision to act (Walton & Bouret, 2019). In addition, our data highlight an important requirement for dopamine acting at NAcC D1Rs in enabling rewards to promote transitions to a vigorous and focused reward-seeking state to allow animals to efficiently achieve their goal. In the former case, activity at NAcC D1Rs increases the likelihood of transitioning to action; in the latter, an absence of activity increases the likelihood of transitioning to an unfocused response state. Therefore, an appropriate balance of activity at NAcC D1Rs is critical to regulate efficient reward seeking. Future studies that employ techniques with greater temporal specificity than is achievable using pharmacology will be helpful to refine this theory.

## Materials and Methods

### Subjects

All procedures were carried out in accordance with the UK Animals (Scientific Procedures) Act (1986). Two cohorts of adult (aged between 8 and 12 weeks at the beginning of training) male Sprague Dawley rats (Harlan, UK) were used in the described studies. Cohort 1 consisted of 11 rats that had previously been implanted for use in an FCV study (Syed et al., 2016), and Cohort 2 consisted of 14 naïve rats. Table 1 outlines which cohorts were used for each pharmacological manipulation, as well as numbers and exclusions for each experiment. Note that the rats in Cohort 1 also received the D1R antagonist systemically, but due to issues with the drug preparation, the incorrect doses were administered and the dataset was excluded. No statistical methods were used to pre-determine sample sizes, but sample sizes are comparable to those reported in previous publications. All animals were maintained on a twelve-hour light/dark cycle. All testing was carried out during the light phase, and during training and testing periods animals were food restricted to 85-90% of their free-feeding weight. Water was provided ad libitum in the home cage.

**Table 1.**
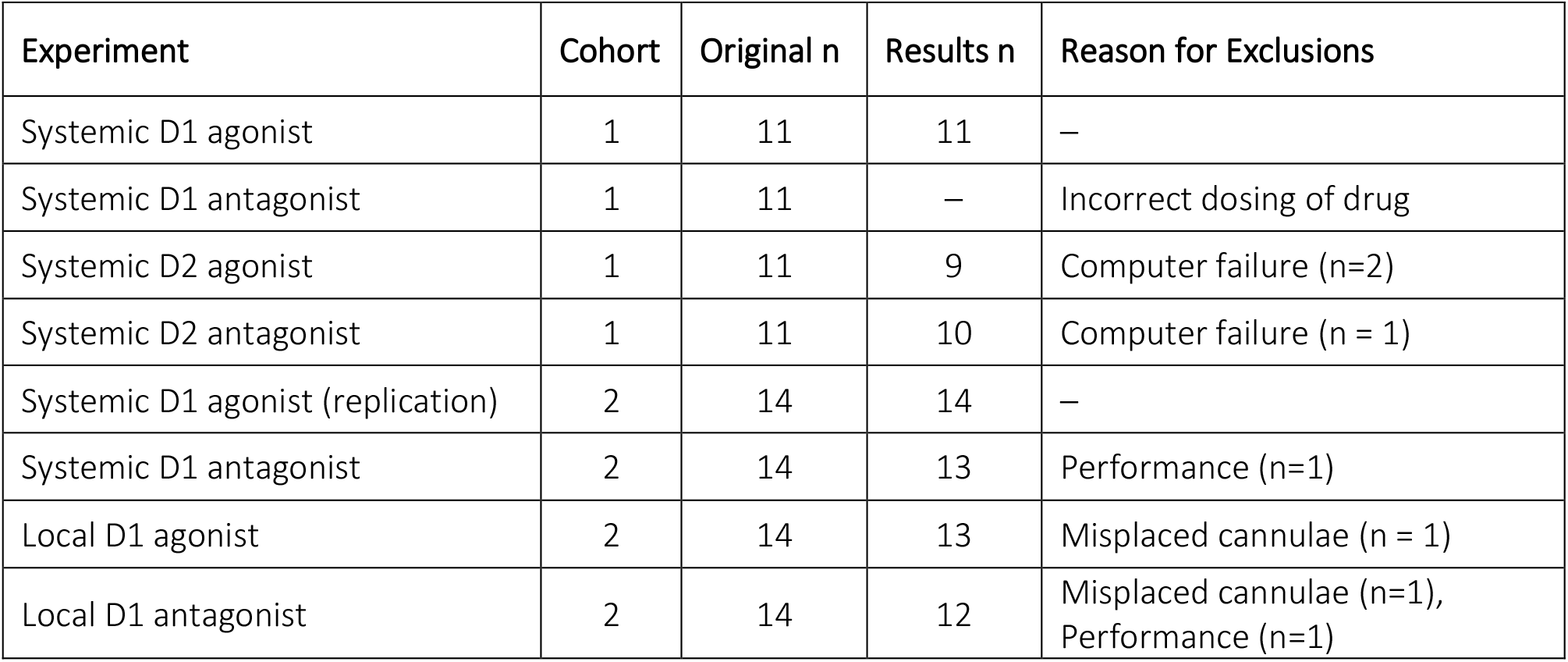
Experimental details, sample sizes and reasons for exclusions. In the case of ‘performance’ exclusions, subjects were excluded if they completed <20% of trials in a session (n = 2 across all experiments).

| Experiment | Cohort | Original n | Results n | Reason for Exclusions |
| --- | --- | --- | --- | --- |
| Systemic D1 agonist | 1 | 11 | 11 | – |
| Systemic D1 antagonist | 1 | 11 | – | Incorrect dosing of drug |
| Systemic D2 agonist | 1 | 11 | 9 | Computer failure (n=2) |
| Systemic D2 antagonist | 1 | 11 | 10 | Computer failure (n = 1) |
| Systemic D1 agonist (replication) | 2 | 14 | 14 | – |
| Systemic D1 antagonist | 2 | 14 | 13 | Performance (n=1) |
| Local D1 agonist | 2 | 14 | 13 | Misplaced cannulae (n = 1) |
| Local D1 antagonist | 2 | 14 | 12 | Misplaced cannulae (n=1),<br>Performance (n=1) |

### Apparatus and behavioural training

Animals were trained on an operant Go/No-Go task (Fig. 1a, b) in which, after initiating a trial by making a nosepoke, auditory cues instructed them either to make (Go) or withhold (No-Go) action in order to gain either a small or large reward. Experiments were conducted using MED-PC behavioural chambers fitted on one wall with two retractable levers 9.5cm on either side of a central nosepoke, and a food magazine on the opposite wall into which 45mg sucrose pellets (Test Diet, Sandown Scientific, UK) were dispensed. Both the nosepoke and food magazine were fitted with infrared beams for entry detection. Each chamber was also fitted with a speaker for delivering the 4 auditory stimuli (∼70dB tone, buzz, white noise, or clicker) and a house light.

After magazine training, animals were first trained on the No-Go trial type. On these trials, the rat was required to remain in the nosepoke for the required period. The No-Go duration was incrementally increased across training sessions on reaching the behavioural criterion (≥ 60% success rate) up to a jittered pre-cue period of 0.3-0.7s and a maximum cued hold period of 1.5-1.7s. A 0.1s ‘buffer’ period was also introduced to distinguish between genuine nosepoke exists and small shifts in posture that may have inactivated the poke detector. Successful trials were rewarded with either one (small reward) or two (large reward) sucrose pellets, as cued by the auditory stimulus.

After reaching criterion for both No-Go trial types, animals were next trained on the Go trial type. Mirroring the No-Go trials, correct choice of one lever (either left or right, side counterbalanced across animals) was rewarded with one pellet (small reward), whilst the other was rewarded with two pellets (large reward), again cued by the auditory stimulus. After again reaching criterion, No-Go trials were interleaved with Go trials to give the full task, in which animals experienced all four trial types pseudorandomly.

### Behavioural task

In the full Go/No-Go task (Fig. 1a, b), animals initiated a trial by entering and remaining in the nosepoke for the pre-cue hold period (0.3-0.7s). This then resulted in presentation of one of 4 auditory cues that indicated the action required (Go or No-Go) and the size of the reward on offer (small or large). Cues were counterbalanced across animals. In No-Go trials, the cue sounded until the end of the hold period or, if the rats exited the nosepoke prematurely, until the time of exiting the poke. In Go trials, the cue sounded until animals pressed the correct lever twice, or until they pressed the wrong lever, or for a maximum of 5s if they failed to press any lever (response omissions).

On correct trials, rewards were delivered 1s after successful completion of a trial (remaining in the nosepoke for the No-Go period on No-Go trials or making 2 correct lever presses on Go trials). After reward delivery a 5s inter-trial interval (ITI) commenced. No cue indicated the end of the ITI and animals were free to initiate the next trial after this time. Both failed Go and No-Go trials resulted in the house light illuminating for a 5s time-out period 1s after the error before turning off and the 5s ITI commencing. The session ended after animals had either gained 100 rewards or after 60 minutes.

### Behavioural measures

All measures were calculated on a session-by-session basis. Performance in this study for each trial type was expressed as percentage success over all attempted trials within a session. On Go trials, animals could make an error by either selecting the incorrect lever (‘WRONG LEVER’), or by omitting responding (‘RESPONSE OMISSION’). On No-Go trials, animals could only make an error by exiting the nosepoke before the end of the cued holding period (‘PREMATURE EXIT’). We reasoned that such premature responses could result from either a failure to inhibit fast cue-driven responses, or from a failure to wait for the appropriate time period before initiating a response. As the former would result in premature responses clustered near cue presentation, and the latter in failures near the end of the holding period, we chose to separately quantify these errors as those occurring in the first (‘EARLY’, < 800ms) or second (‘LATE’, > 800ms) half of the No-Go holding period. This was calculated as a proportion of all No-Go nosepoke exits in that session. As an additional metric of impulsive responding, we also measured the number of head exits made during the pre-cue period (after a nosepoke was made to initiate a trial, but before a cue was presented), which were termed ABORTED trials.

Key task latencies in all successful trials included: (a) **ACTION INITIATION:** cue onset to nosepoke exit (NB. on Go trials, this did not include trials in which animals remained in the nosepoke >1.7s, i.e. indicating that the Go trial was interpreted as No-Go trial), (b) **TRAVEL TIME (Go trials only):** time from nosepoke exit to first lever press; and (c) **REWARD RETRIEVAL:** time from reward delivery to magazine entry. Additionally, we calculated (d) **RE-ENGAGEMENT:** latency from magazine entry to re-entering the nosepoke (after a successful trial, and regardless of whether this was during or after the ITI).

### Video tracking

Videos were captured at 25 fps and video tracking was performed using the DeepLabCut toolbox (Mathis et al., 2018). Two separate models with matching parameters were trained to account for differences in box orientation and nodes included the rats’ nose, ears, head, body, legs, and tail, as well as key features of the operant chambers – the nosepoke, left and right levers, and left and right corners of the food magazine. For each video, 25 randomly selected frames were manually labelled before the network was trained and tested over 1030000 iterations, resulting in an average tracking error of < 5 pixels. After training, only frames with co-ordinates that had a likelihood of 1 were included and any missed frames were interpolated across. These co-ordinates were aligned with MED-PC behaviour data by identifying when animals made errors in the task; errors resulted in the houselight being turned on such that the average luminance values from greyscale converted video frames increased sharply, and were therefore identifiable using the inbuilt MATLAB ‘findpeaks’ function. As the camera system had not originally been set up with the intention of performing such granular analyses, a number of sessions had to be excluded due to suboptimal video quality. Reasons for excluding a session included a failure to align tracking with MED-PC behaviour, poor visibility of operant chamber features, and sessions in which a majority of frames required interpolation. X and y co-ordinates from the tracked nose marker were used for all analyses. Any negative values in the y axis were due to the rat having its nose in the nosepoke and were therefore converted to 0.

### Video analyses

We normalised the values of operant chamber parts on a per session basis by subtracting the median co-ordinates of the nosepoke from the values of the tracked parts. For all behavioural metrics relating to analysis of rat location in the chamber relative to chamber components (aside from the calculation of entropy – explained below), we first divided the range of box space co-ordinates into a 3 x 3 grid, allowing for two squares of the grid to be labelled as lever squares, one as a nosepoke square, and one as a magazine square. For each frame, if a given co-ordinate was within the boundaries of a square it was scored as ‘1’, or ‘0’ otherwise. This was averaged across to give a mean probability density function, or probability, across trials for each rat, and then averaged across rats. To calculate the time spent in the area of the nosepoke we measured the latency from the beginning of the trial to when the animals were detected to have left this square based on the tracked nose co-ordinates being outside of the boundary of the nosepoke square. To calculate the proportion of trials on which animals completed a particular sequence of actions, e.g. moving from the correct lever to the food magazine, we again used this binary measure of whether the tracked co-ordinates had entered within the specified squares but in addition specified a required order of visitation for the trial to be counted as such. Trajectory lengths were calculated by finding the Euclidean distance between frames and normalised by the median distance between the lever markers before being summed and averaged. For the calculation of entropy, we divided the range of box space co-ordinates into an 18 x 18 grid in order to increase the granularity of the analysis. We then normalised the probability values of each location, *L*, in the grid on a per session basis before calculating entropy using the Shannon equation (Shannon, 1948) for each session as: ***E* = − ∑ *p***_***L***_ ***log***_**2**_***p***_***L***_. For the colourplot visualisations in Fig. 6 and Fig. 8, the box boundaries were reduced by 50 pixels along the x axis as very few tracking points reached those co-ordinates.

### Pharmacological challenges

SKF 81297 hydrobromide (Tocris Bioscience; D1R agonist) was administered systemically at doses of 0.3 mg/kg and 1.0 mg/kg, and locally at doses of 0.4 μg/μl and 4.0 μg/μl. SCH 23390 hydrochloride (Tocris Bioscience; D1R antagonist) was administered systemically at doses of 0.005mg/kg and 0.01mg/kg, and locally at doses of 0.2μg/μl and 2.0μg/μl. Quinpirole hydrochloride (Tocris Bioscience; D2R agonist) was administered systemically at doses of 0.0125 mg/kg and 0.0375 mg/kg, and eticlopride hydrochloride (Tocris Bioscience; D2R antagonist) was administered systemically at doses of 0.01 mg/kg and 0.03 mg/kg. Doses were calculated as the salt. All drugs were dissolved in 0.9% sterile saline, made in batch, aliquoted, and frozen at −20°C. Individual aliquots were defrosted for use on testing days. 0.9% sterile saline was administered in control sessions. In all experiments, doses were applied in a counterbalanced Latin square approach, although blinding was not used when making up or applying agents.

### Surgical procedure

To implant cannulae targeting the NAcC, rats were anaesthetised using inhaled isoflurane (4% vol/vol in O_2_ induction and 1.5% for maintenance delivered via facemask) and administered buprenorphine (Vetergesic, 0.03 mg/kg, s.c.), meloxicam (Metacam, 2mg/kg, s.c.) and 3ml glucosaline (Aquapharm). Body temperature was maintained at 37±0.5°C by a homeothermic heating blanket. Once animals were secured in a stereotaxic frame (Kopf Instruments) and their scalp shaved and cleaned with dilute hibiscrub and 70% alcohol, a local anaesthetic (bupivacaine, 2mg/kg) was administered to the incision site. Eye gel (Lacri-Lube, Allergan) was applied to the eyes for protection. The skull was then exposed and the skull was levelled based on measurements of Bregma and Lambda. Six holes were drilled: two for implantation of bilateral guide cannulae (Plastics One, UK) and four for anchoring screws (Precision Technology Supplies). Guide cannulae were then lowered. The cannulae consisted of an 8mm plastic pedestal holding two 26-gauge metal tubes with a centre-to-centre distance of 3.4mm and a length of 7.5mm. They were implanted 1.5mm above the target site of the NAcC, at co-ordinates of AP relative to bregma: +1.4mm, ML: ±1.7mm, DV: - 6.0mm from surface of skull relative to bregma. Dental acrylic (Associated Dental Products Ltd.) was then applied to secure the cannulae to the skull and screws. After surgery, bilateral dummy cannulae were inserted to ensure patency, and a dustcap was secured to the pedestal. Animals were again administered buprenorphine, meloxicam, and glucosaline post-surgery, and meloxicam was given up to a further three days post-surgery. Animals were group housed after initial recovery and began re-training once they were fully recovered, on average two weeks after surgery.

### Systemic administration procedure

Drugs were injected intraperitoneally (i.p.) at a volume of 1 ml/kg bodyweight. All drugs were injected 10 minutes before the behavioural session aside from the D1R antagonist, which was administered 20 minutes before the start of the session. Drug administration sessions were separated by at least one treatment-free training day to ensure a return to baseline performance and complete washout of the drug, with criteria of ≥ 60% successful trials across all trial types, and session completion within 60 minutes. If these criteria were not met, animals continued with treatment-free training days until performance reach criteria, at which point the next testing day commenced, though in practice animals’ performance almost always reached criteria at the first training day.

### Local administration procedure

A mock infusion was carried out one day prior to the first experimental session to reduce potential tissue damage-related confounds. This involved insertion of the injectors back-filled with saline without infusion of any substance. The following day two 10μl glass Hamilton syringes were back-filled with 0.9% sterile saline and placed in an infusion pump (Cole-Parmer). Double connector assembly tubing (Plastics One, UK) was cleaned with ethanol and thoroughly dried with air, then filled with saline before being attached to the Hamilton syringes. 33-gauge 9mm bilateral injectors (Plastics One, UK) that had been cleaned by sonication for one hour in 70% ethanol were then attached to the connector assembly. A small air bubble separated the saline from drug. The injectors were checked for blockage before the rats were gently restrained, their dummy cannulae removed, and injectors inserted. During infusion, 0.5μl of solution was injected per hemisphere at 0.25μl/minute. Injectors were left in place for a further two minutes after infusion and then removed. The dummy cannulae and dustcap were replaced and rats were returned to their homecage for 10 minutes before beginning the task.

### Data analysis

All datasets are available from the corresponding authors on reasonable request. Data was extracted and analysed using MATLAB R2018a and IBM SPSS Statistics 24. Significant interactions were explored by analysis of the simple effects and are reported in the appropriate figure legends or tables. Randomisation or blinding was not used during analysis.

### Histology

At the end of data collection, cannulated animals were deeply anaesthetised with sodium pentobarbitone (200mg/kg, i.p. injection) and transcardially perfused with 0.9% saline followed by a 10% formalin solution (vol/vol). Brains were kept in 10% formalin solution until being sectioned. Brains were sectioned into 60μm-thick coronal sections by vibratome (Leica). The sections were stained with cresyl violet (Sigma Aldrich) before being mounted in DePeX mounting medium onto 1.5% gelatin-coated slides and enclosed with coverslips to confirm cannulae placements (Supp. Fig. 4).

## Acknowledgements

We would like to thank Greg Daubney for help with histology. This work was supported by Wellcome (fellowships WT090051MA and 202831/Z/16/Z to MEW, 206330/Z/17/Z to MH), the ESRC (award ES/J500112/1 to LLG) and the Clarendon and the Archimedes Foundation (awards SFF1819_CB2_MSD_1196514 and Kristjan Jaagu scholarship to OH). The authors declare no competing financial interests. For the purpose of open access, the author has applied a CC BY public copyright license to any Author Accepted Manuscript version arising from this submission.

## Author Contributions

LLG, MH and MEW conceived the project. LLG, ES and OH trained the animals. LLG performed the surgeries with assistance from MCP and OH. LLG collected the data, with assistance from MCP for the local infusion studies. LLG and MEW analysed the data with input from MCP and SGM. LLG and MEW prepared the manuscript with input from all the other authors.

## Supplementary text

### *Supp. Text 1:* Global D2R stimulation modulates action vigour

To understand how specific the observed systemic effects were to D1R manipulation, we also investigated the effect of systemic administration of either a D2R agonist or antagonist (both cohort 1). The D2R agonist increased the proportion of premature responses during the pre-cue period (main effect of drug: *F*_(2,16)_ = 3.652, *p* = .049), but neither the agonist nor antagonist had any effect on No-Go success, time spent in the nosepoke on successful No-Go trials, nor the distribution of early and late No-Go errors (all *p* > .1, data not shown).

However, the D2R agonist did markedly slow reward retrieval latencies in No-Go trials (main effect of drug: *F*_(2,14)_ = 23.044, *p* < .001) and this effect was paralleled by an overall slowing of movements in Go trials. It strongly decreased success rates (Supp. Fig. 1a; main effect of drug: *F*_(2,16)_ = 45.299, *p* < .001) – mainly due to an increase in response omissions (Supp. Fig. 1b; main effect of drug: *F*_(2,16)_ = 58.576, *p* < .001; lever selection errors, Supp. Fig. 1c; all *p* > .06) – and also slowed all latencies, including action initiation (at the highest dose), travel time, and reward retrieval (Supp. Fig. 1d-f; all *F* > 7, *p* < .004; also reward retrieval: drug x reward interaction: *F*_(2,16)_ = 7.962, *p* = .005). In contrast, the D2R antagonist had no reliable effect on any measure of either No-Go or Go performance or response time (Supp. Fig. 1g-l; all *p* > .08). Together, these results show that stimulating D2Rs reduces the vigour of all actions – both in Go and No-Go trials – an effect distinct to the influence of D1Rs which mainly affected the vigour of actions distal to reward.

**Supp. Fig. 1.**
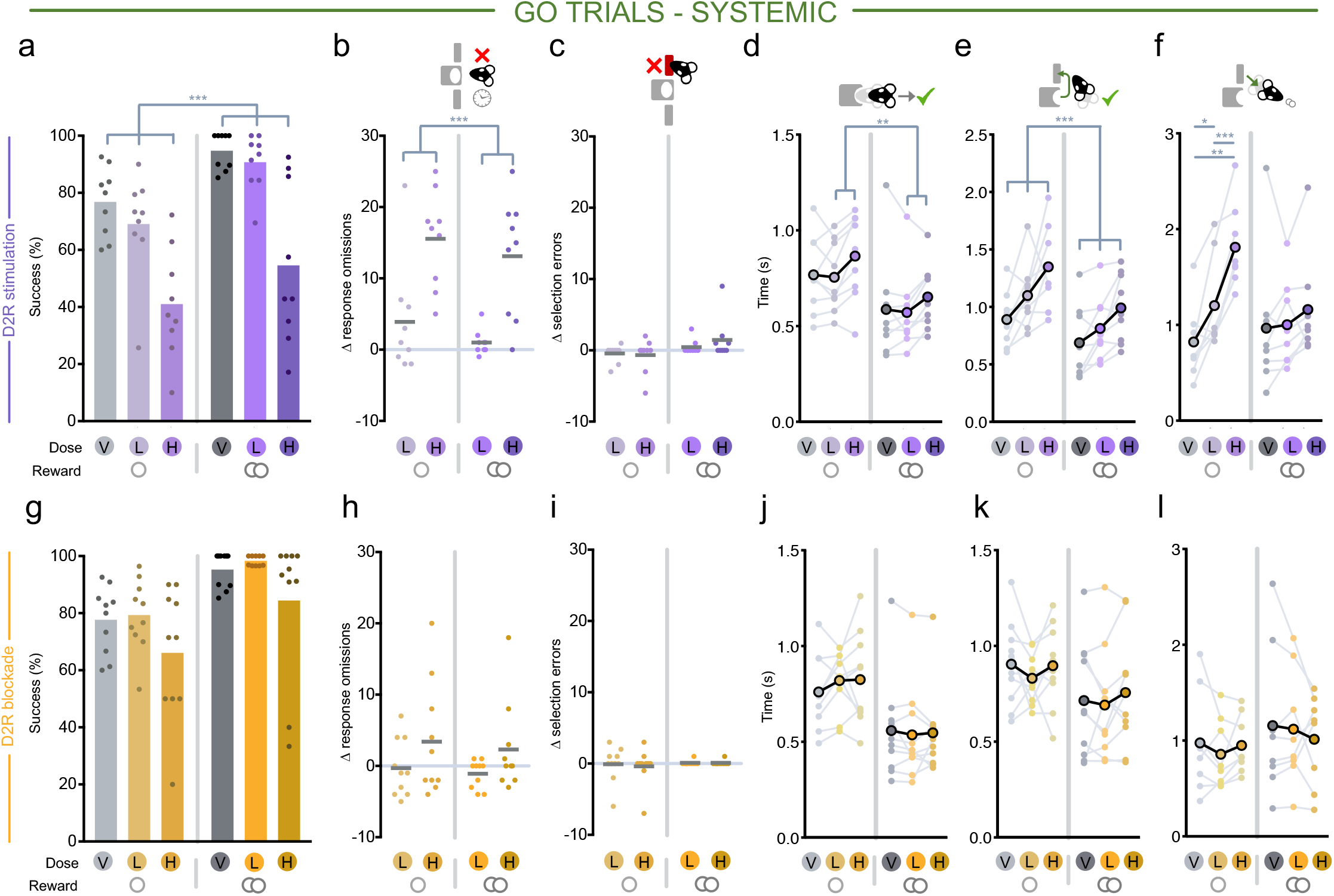
Systemic effects of D2R stimulation (quinpirole) or blockade (eticlopride) in Go trials. V = vehicle, L = low dose, H = high dose. Single circle indicates small reward condition, double circle indicates large reward condition. **(a-f)** Effects of D2R stimulation split by small (left) and large (right) reward Go trials on **(a)** success rate, **(b)** response omission errors, **(c)** lever selection errors, **(d)** latency to leave the nosepoke after Go cue onset, **(e)** latency from nosepoke exit to first lever press, **(f)** and latency from trial completion to entering the food magazine to retrieve reward. **(g-l)** Same as in **(a-f)** but for systemic D2R blockade. \*\*\**p* < .001, \*\**p* < .01, \**p* < .05

**Supp. Fig. 2.**
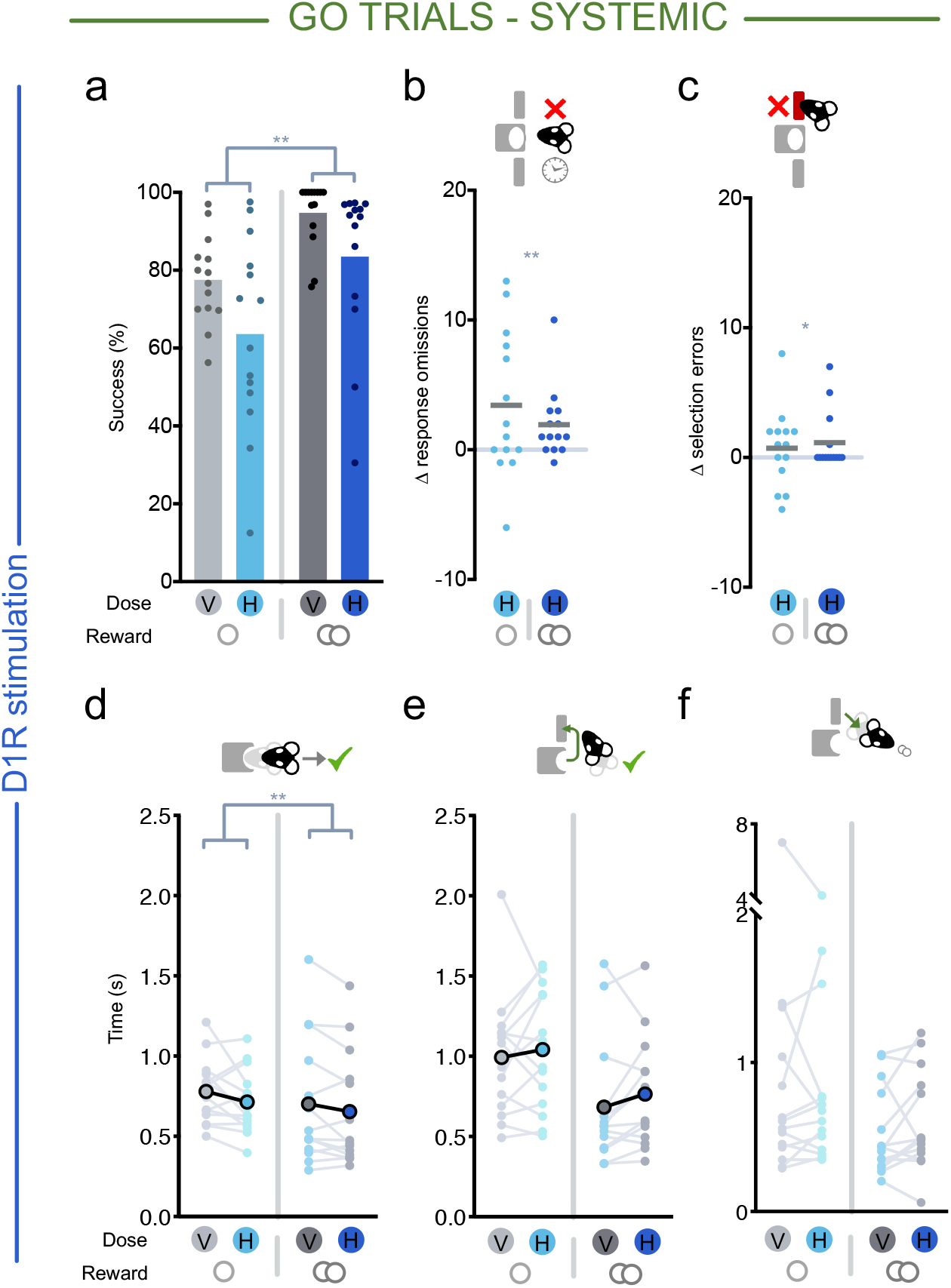
Systemic D1R stimulation replication study results. Effect of D1R stimulation on **(a)** success rate (main effect of drug: *F*_(1,13)_ = 14.237, *p* = .002), **(b)** response omission errors (main effect of drug: *F*_(1,13)_ = 11.424, *p* = .005), **(c)** lever selection errors (main effect of drug: *F*_(1,13)_ = 4.694, *p* = .049) **(d)** latency to leave the nosepoke after Go cue onset (main effect of drug: *F*_(1,13)_ = 10.895, *p* = .006), **(e)** latency from nosepoke exit to first lever press (main effect of drug and interaction n.s., *p* > .1), **(f)** and latency from reward delivery to entering the food magazine to retrieve reward (main effect of drug and interaction n.s., *p* > .3). ** *p* < .01, * *p* < .05

**Supp. Fig. 3.**
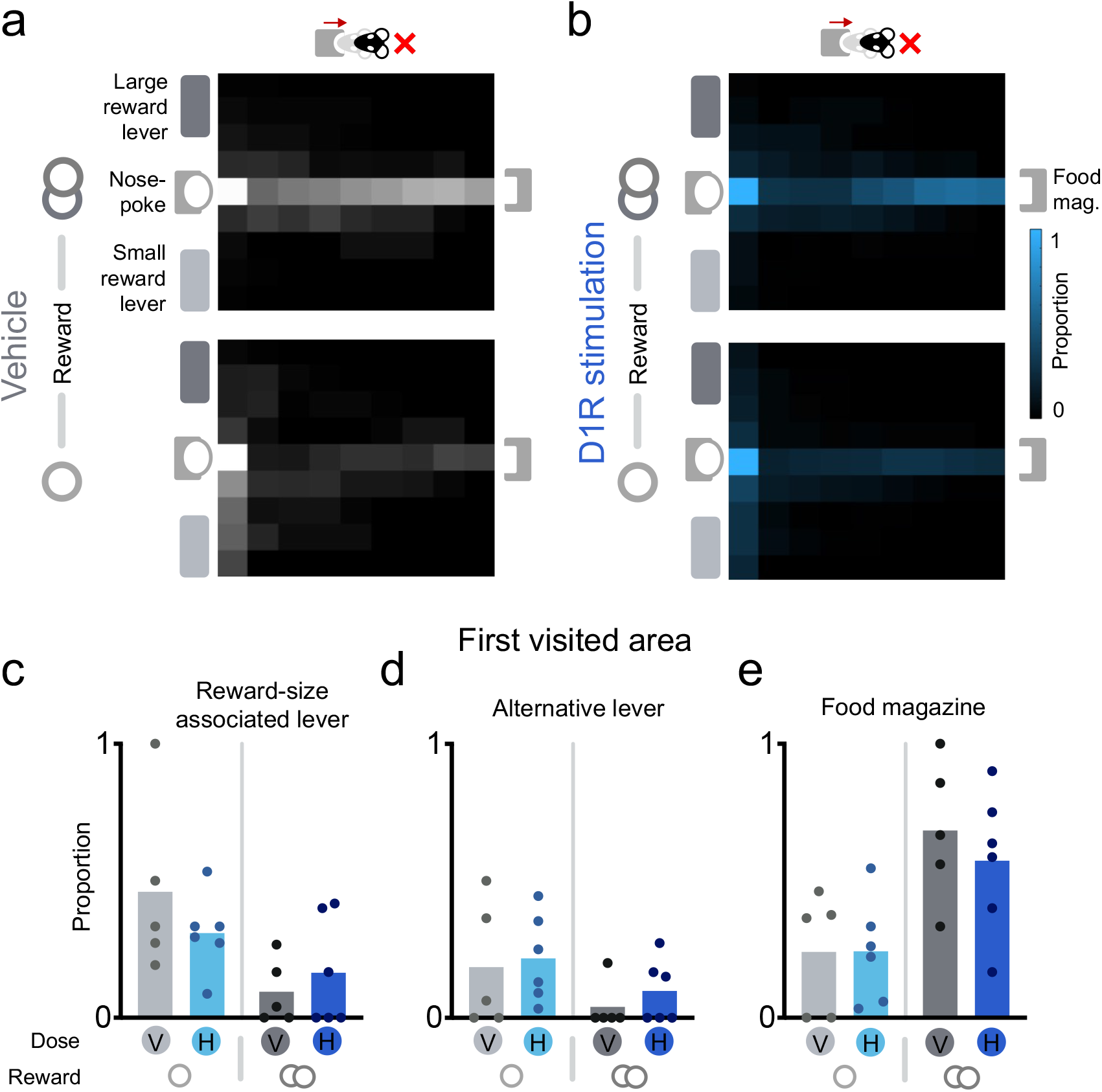
The effects of intra-NAcC D1R stimulation (SKF-81297) in error No-Go trials. **(a, b)** Mean probability density across rats in small (lower) or large (upper) reward error No-Go trials when **(a)** on vehicle or **(b)** with intra-NAcC infusion of the D1R agonist. **(c-e)** Proportion of trials in which the first area of the operant chamber visited by the rats was **(c)** the reward size-associated lever, corresponding to the large reward lever on large reward No-Go trials and the small reward lever on small reward No-Go trials, **(d)** the alternative lever, and **(e)** the food magazine. Location pairwise comparisons: reward-size associated lever vs. food magazine: *p* = .032, alternative lever vs. food magazine: *p* = .002, reward-size associated lever vs. alternative lever: *p* = .196.

**Supp. Fig. 4.**
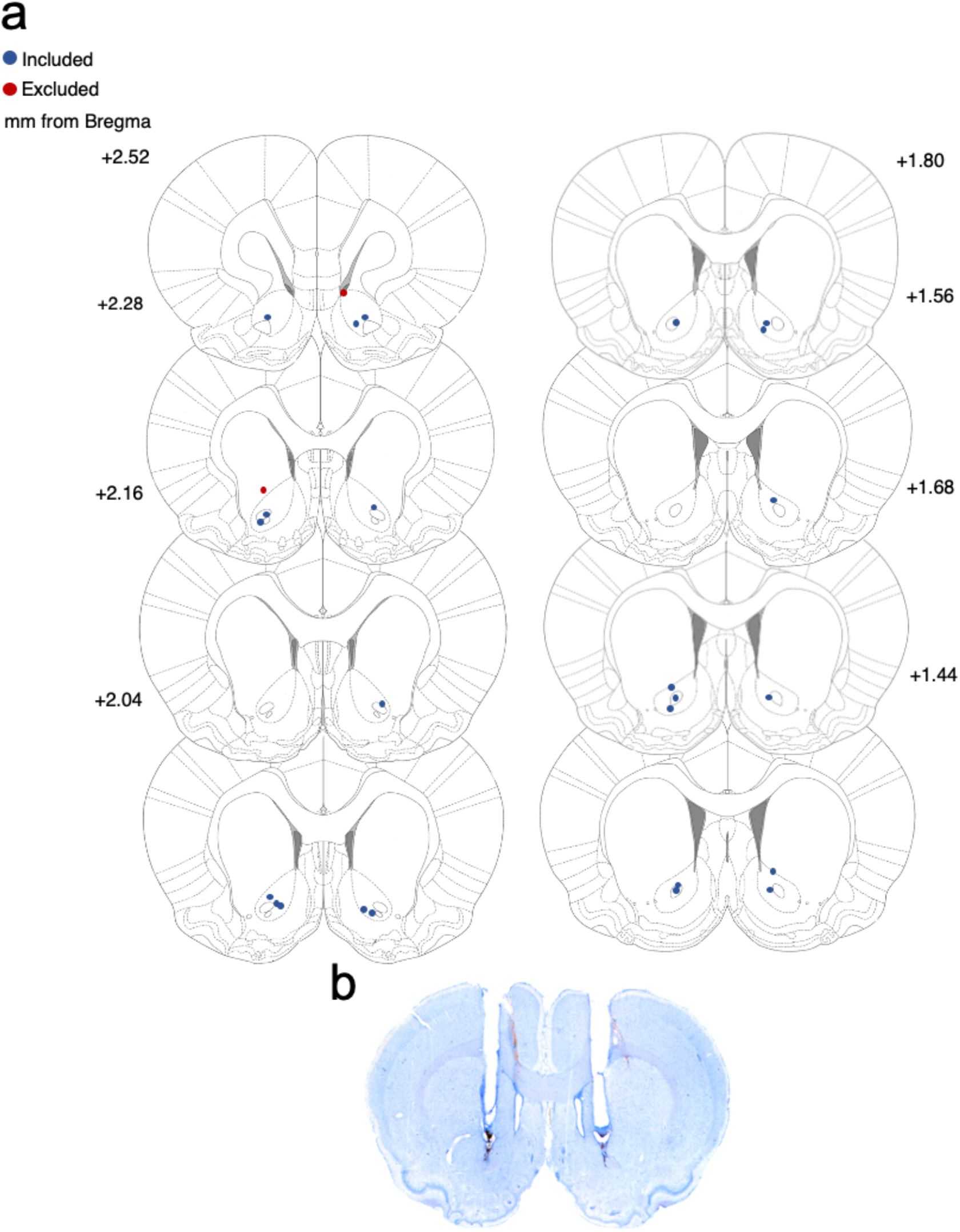
Injector placement. **(a)** Schematic of cannula insertion locations in the NAcC (n = 14, all rats bilaterally implanted). Cannulae locations of included rats are marked in blue, excluded rats (n = 1) are marked in red. Numbers to the left of coronal sections indicate distance anterior to bregma (mm). Adapted from the atlas of Paxinos and Watson (2009). **(b)** Example photo scan of a perfused section showing bilateral injector lesion and guide cannulae placement.

